# Conserved sequence elements in the final exon of MDM2-eight-exon skipping event reveal a ‘cassette regulon’ model of alternative splicing controlled by a distal regulatory element

**DOI:** 10.64898/2026.02.14.705925

**Authors:** Safiya Khurshid, Matias Montes, Rafia Rahat, Zac LaRocca-Stravalle, Whitney Jimenez, Andrew Goodwin, Claudia Kelly, Hannah Ackerman, Brad Bolon, Vincenzo Coppola, Dawn S. Chandler

**Author notes:** Corresponding author: Dawn S. Chandler, PhD. Equal contribution (Safiya Khurshid, Matias Montes).

## Abstract

Modern sequencing technologies have revealed that cancers exhibit widespread dysregulation of alternative splicing, which plays a significant role in driving tumor hallmarks. Alternative splicing enables a single gene to generate multiple mRNA variants, a fundamental mechanism that shapes the distinctive characteristics of normal cells as well as cancer cells. These splicing events are governed by complex interactions between *cis*- and *trans*-acting factors and understanding these mechanisms is crucial for effectively targeting cancer cells. This study highlights a unique splicing event in which genotoxic stress induces the skipping of eight-exons in the proto-oncogene *MDM2*, resulting in the formation of *MDM2-ALT1*, an isoform that is overexpressed in various cancers. To effectively target this splicing event, it is important to unfold the regulatory mechanism behind it. We hypothesize that the event occurs either by the autonomous regulation of individual exons (*Exon autonomous model*), where each exon is regulated independently or by the coordinated exclusion of an eight-exon-regulon unit (*Exon Regulon model*). Utilizing *in-silico* tools, a comprehensive modular minigene system, CRISPR-mutated cell line, and murine models, we demonstrate that this complex *MDM2* splicing event is regulated by sequences within a distal terminal exon, which emerge as critical regulators, orchestrating the alternative splicing of *MDM2*. Constitutive expression of the *Mdm2-MS2* isoform (the mouse ortholog of the human *MDM2-ALT1* splice variant), achieved by mutating the *Srsf2* binding-site on exon 11, modulates proliferation and apoptosis dynamics in NIH3T3 cells. Moreover, in a p53-wildtype *in-vivo* setting, this isoform confers a protective effect against age-induced neoplasia. Our findings support the *Exon Regulon model* of *MDM2* splicing, regulated by distal elements analogous to distal enhancer elements that control transcription. These finding sheds light on intricacies in the splicing code that could have significant implications for developing splice variant targeting cancer therapies.

## Introduction

Alternative splicing represents a widespread regulatory process in gene expression, that enables the production of multiple distinct mRNA variants from a single gene (1). Understanding the detailed mechanisms of this process is crucial, as alternative splicing drives many defining characteristics of cancer cells (2). Cancer-related mutations in splicing factors or changes in their expression levels can disrupt normal splicing patterns of key regulatory genes involved in cell proliferation, apoptosis, and metastasis by generating oncogenic isoforms that disrupt tumor suppressor functions (3,4). The regulation of alternative splicing is orchestrated through a complex network of *cis*- and *trans*-acting factors, and is further influenced by the dynamic interplay between transcription and splicing processes (5). The inclusion or exclusion of specific exons, as well as intron retention, in mature mRNA is shaped by intrinsic features such as splice-site strength, exon length, and conserved splice-site sequences surrounding orthologous alternative exons, which together create a regulatory landscape that can be modulated by external stimuli such as cellular or genotoxic stress (5–8). In cancer cells, these regulatory mechanisms are often exploited to generate splice variants that provide the tumor with advantages, such as increased proliferation, immune evasion, or resistance to therapy. Therefore, elucidating the molecular basis of alternative splicing is crucial for developing effective strategies to target cancer cells (9).

Several *trans*-acting factors influence splicing, with the serine-arginine-rich (SR) class of proteins being among the most well-studied regulators (8,10,11). These proteins promote or repress the inclusion of specific exons in nascent pre-mRNA transcripts (10,12,13). Maintaining balanced levels of these factors and their interaction with designated pre-mRNA sites, whether under homeostatic conditions or in response to external stimuli, significantly influences spliceosome assembly and splice-site selection. Serine/arginine-rich splicing factor 2 (SRSF2), also known as SC35, like its other family members is characterized by an RNA recognition motif (RRM) and a arginine/serine-rich (RS) domain. SRSF2 regulates pre-mRNA splicing by binding to specific sites on the pre-mRNA through its RRM motif and interacting with spliceosome components or other splicing factors through its RS domain (14–16). SRSF2 is a well-known positive regulator of splicing. Its involvement in processes such as the regulation of exon 10 in *tau* and the splicing of *CD44* pre-mRNA (17,18) reinforces its canonical role in promoting exon inclusion.

Murine Double Minute 2 (*MDM2*) functions as a proto-oncogene, exerting critical control over the tumor suppressor protein, p53. The full-length *MDM2* transcript comprises 12 exons and encodes a protein with a p53 binding domain and an E3 ubiquitin ligase domain, which facilitates p53 ubiquitination and subsequent degradation (19–21). Under genotoxic stress (e.g. UV-irradiation or cisplatin treatment), *MDM2* undergoes alternative splicing, to produce a spliced variant known as *MDM2-ALT1(MDM2-B*), which retains only coding exons 3 and 12 and excludes exons 4 through 11 (22,23). This multi-exon splicing event is observed in several cancers and is conserved in murine cells, revealing evolutionary conservation of the *MDM2* regulated splicing. In mice, the analogous splicing event yields the isoform *Mdm2-MS2* (containing exons 3, 4 and 12), a mouse ortholog of human *MDM2-ALT1* (7). Previous work from our laboratory has established that SRSF2 acts as a positive splicing factor, facilitating the inclusion of exon 11 in *MDM2* and thereby promoting the production of full-length *MDM2*. Disruption of the SRSF2 binding-site within exon 11 results in the exclusion of this exon from the mature transcript (24). However, the mechanism by which the complex multi-exon skipping event is coordinated in the presence of genotoxic stress and/or in an oncogenic background is not clear. We considered two possible hypothesis for regulation of *MDM2:* 1. Each exon (4 through 11) is regulated independently, a scenario we term the *’exon autonomous model* ’. 2. Exons (4 through 11) could be regulated collectively as a single unit, which we refer to as the ’*exon regulon model’*. Given that *MDM2* undergoes alternative splicing in cancers and is associated with high-grade disease (25), understanding the regulatory mechanism of this splicing event will inform the development of improved treatment approaches. Our research presented here reveals that exon 11 is pivotal in coordinating this splicing event and governs the splicing of its preceding seven exons. This research deepens our understanding of splicing regulation and its functional outcomes, offering novel insights into the distal elements that control splicing decisions demonstrating functional parallels to distal enhancers that control transcription.

## Results

### MDM2 minigene panel reveals a lack of DNA damage–induced splicing regulation in exons 4–10

*MDM2* is an important inhibitor of the tumor suppressor protein TP53, and its splicing is frequently deregulated in a number of cancers (25–27). The most prominent tumor-associated isoform, *MDM2-ALT1* (also known as *MDM2-B*) is characterized by the direct splicing of exon 3 to exon 12, resulting in the exclusion of eight-internal exons that encode the p53-binding domain, ARF-binding domain and the nuclear localization signal (Figure 1A). To effectively target this splicing event in cancer, a comprehensive understanding of the mechanistic regulation of *MDM2* alternative splicing is crucial.

**Figure 1:**
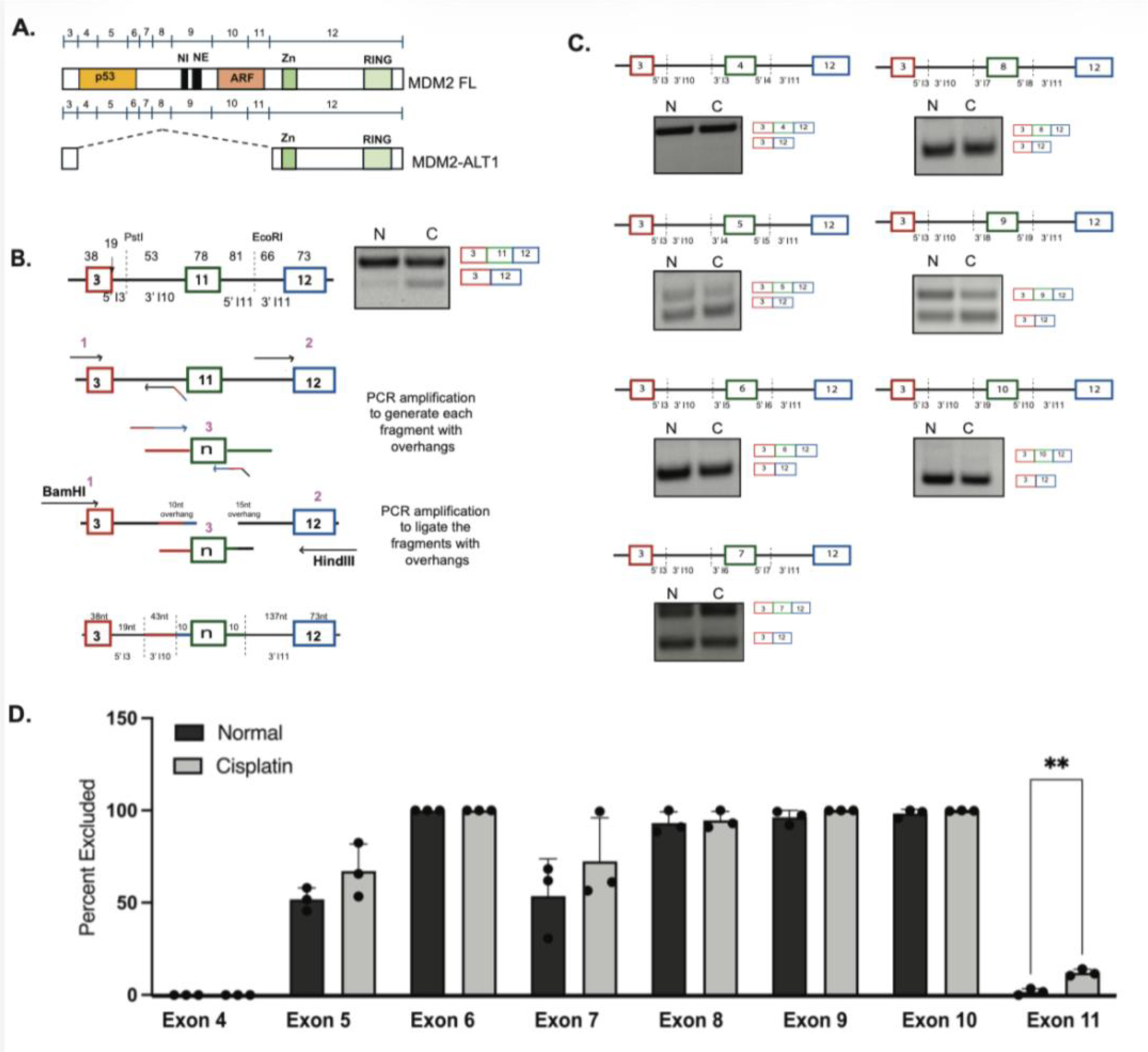
*MDM2* minigene panel reveals a lack of DNA damage–induced splicing regulation in exons 4–10. **A.** Schematic representation of alternative isoforms of *MDM2*. **B**. Schematic representation of modular minigene construction. Parent 3-11-12 minigene was used and exon 11 was swap for other exons that are shown. **C**. Hela cells were transfected with respective minigenes, and cells were treated with 75uM cisplatin and harvested after 24-hours. RT-PCR gels representing minigene splicing before and after damage using *MDM2* minigene primers (Primer information in Supplementary Table 5). **D**. A bar graph showing the quantification of n=3 experiments. **E.** Summary table representing the damage inducible splicing status of individual exons.

To explore the mechanistic role of *cis-* and *trans-*regulatory elements in this process, we initiated an *in-silico* analysis focused on the splicing factor SRSF2, previously identified in our lab as a key regulator of damage-inducible splicing in *MDM2* (24). We investigated whether SRSF2 preferentially binds to specific exons, potentially influencing their likelihood of being skipped under genotoxic stress conditions. Using ESE finder (6, 26), we analyzed each *MDM2* exon for SRSF2 binding motifs (Supplementary Table 1). The binding-site score revealed that each exon contains one or more predicted SRSF2 binding sites, and that all but one exon (Exon 7) has SRSF2-sites that are conserved in mice (either by position within the exon, or sequence or both). This suggests that each exon may be individually regulated by SRSF2, and that *MDM2* splicing could follow an ‘*exon autonomous model*’ of *MDM2* splicing.

To experimentally test the ability of each exon to be regulated autonomously, we designed a modular *MDM2* minigene panel derived from our laboratory’s pre-existing damage inducible 3-11-12 *MDM2*-minigene (Figure 1B, top panel) (23,29). Using its design framework, we constructed minigenes, in which each individual exon 4-10, accompanied by minimal intronic sequences, is cloned in place of exon 11 (Figure 1B). These *MDM2* minigene constructs were then transfected into HeLa cells using standard lipid-based transfection for 24-hours and treated with cisplatin to induce genotoxic stress for an additional 24-hours. 48-hours post transfection, total RNA was harvested and used to capture the splicing of these minigenes in response to genotoxic stress. RT-PCR data indicates that the minigenes were capable of supporting splicing in 3 different ways: a) All three exons are included (intervening exon 4); b) All three exons are included concomitant with the intervening exon being excluded (intervening exons 5, 7) and c) the intervening exon is consistently excluded (intervening exons 6, 8, 9 and 10) (Figure 1C, 1D, 1E).

However because 4 of our intervening sequences did not support any inclusion of the exon in the final transcript, we hypothesize that these exons are not recognized by the splicing machinery due to the incorporation of only minimal, potentially insufficient intronic sequences flanking the exon. These exons are therefore unable to be further excluded in response to stress. We therefore sought to improve the design of the minigene panel by including additional intronic sequences so as to better assess damage induced splicing regulation.

### Incorporation of longer native 5’ and 3’ intronic sequences rescues exon recognition in the minigenes

Exon recognition is highly dependent on surrounding intronic cues, including native branch points, polypyrimidine tract sequences and splice-site proximal regulatory motifs. Consistent with this requirement for exon recognition, our minigenes containing intervening exons 6, 8, 9 and 10 were likely unable to produce full-length *MDM2* due to insufficient native intronic context. To address this, we performed additional bioinformatic analysis to verify whether the incorporation of increased native intronic regions would change the predicted splice-site strength score. Using bioinformatic tools, such as the Berkeley Drosophila Genome Project’s splice-site predictor and polypyrimidine tract scoring, we derived a 3’ splice-site strength score for *MDM2* introns 3 through 11 (28,30) (Supplementary Table 2). Based on this analysis, we engineered a new set of minigenes with additional native intronic sequences flanking both splice-sites. Notably, the splice-site strength of these new minigenes were restored to those of the endogenous splice-sites. Though, the previous generation of minigenes with minimal intronic sequences consistently maintained their donor site splice-site score, all but exon 11 made weaker acceptor splice-site scores with 3 exons (6, 8 and 9) having no detectable splice-site at all (Supplementary Table 3). These deficiencies in acceptor-site scores could explain why constructs containing intervening exons 6, 8, 9 and 10 failed to produce the full-length transcript.

Having determined through bioinformatic analysis that incorporation of longer native intronic sequences rescued predicted splice-site strength (Supplementary Table 3), we next examined whether any of these new minigenes exhibited damage induced splicing. Following the same protocol as described above, we transfected these new constructs into HeLa cells and treated them with cisplatin to induce genotoxic stress. After treating cells for 24-hours, we harvested RNA and performed RT-PCR to assess the effect on splicing. The data revealed that in addition to original exon 11 minigene construct, constructs with intervening exon 4 and exon 5 now also exhibited damage-inducible skipping (Figure 2A-C). Although minigenes with intervening exons 6, 7, 8, 9 and 10 remained non-damage inducible, the data shows that restoring their native intronic splice-sites promoted exon-recognition by the spliceosome, resulting in exon inclusion across all these minigenes constructs (Figure 2B).

**Figure 2:**
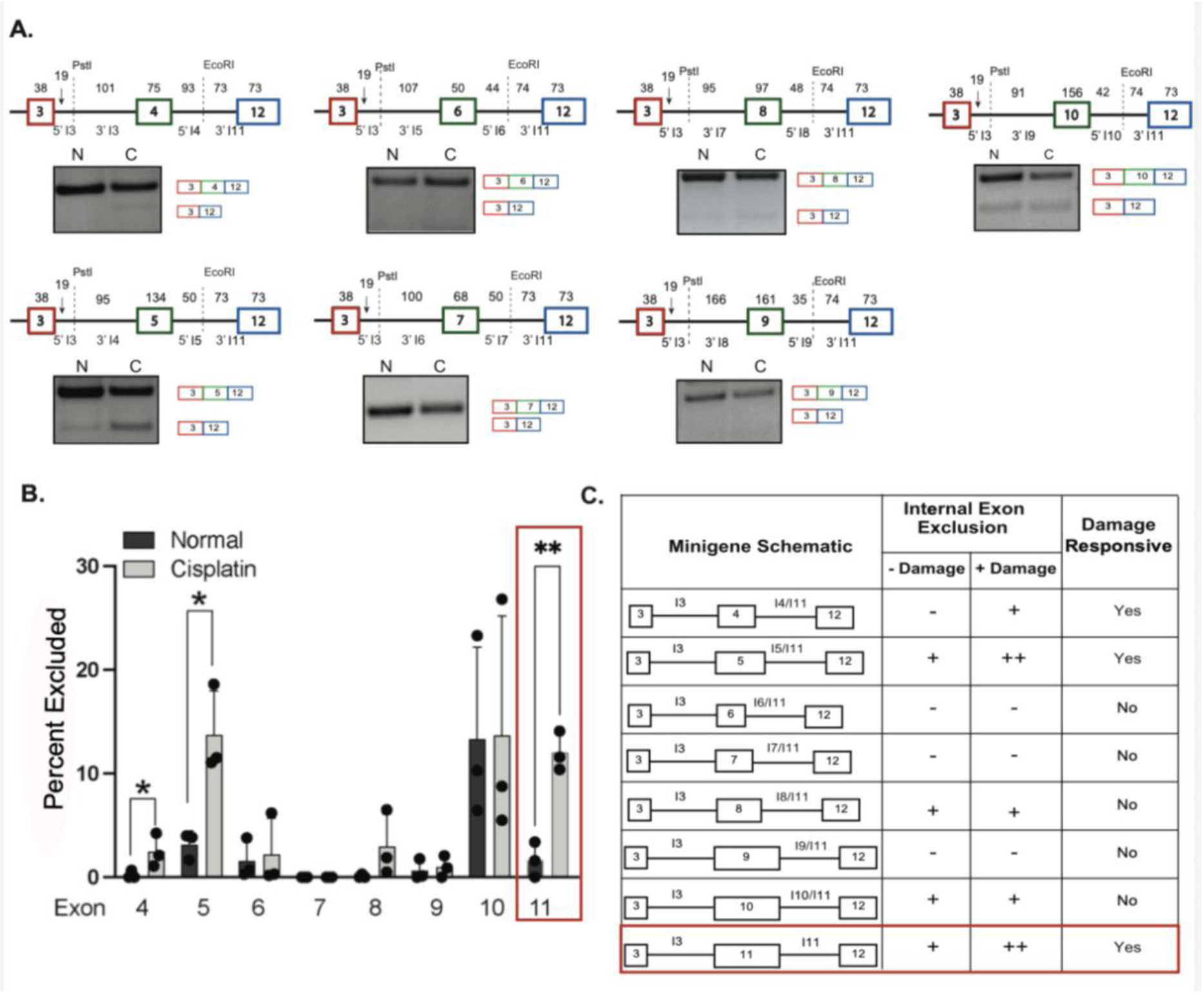
Incorporation of longer native 5’ and 3’ intronic sequences rescues exon recognition in the minigenes. **A**. Hela cells were transfected with newly constructed minigene set II, treated with cisplatin and harvested after 24-hours. The gel images represent RT-PCR using *MDM2* minigene primers. **B**. Bar graph showing the quantification of n=3 independent experiments. Data from Figure 1 for Exon 11 (in red box) is shown for comparison. **C.** Summary table representing the damage inducible splicing status of each individual exon when native intronic sequences were reincorporated.

This rescue in exon-recognition combined with the selective occurrence of damage-induced skipping, observed in exons 4, 5 and 11 but not exons 6-10 highlights an outcome that is exon dependent . Given our initial bioinformatic analysis on predicted SRSF2 binding sites in all *MDM2* exons, these variable splicing outcomes prompted us to closely examine the SRSF2 binding scores for each exon. To identify the most functionally relevant SRSF2 predicted binding-sites, we adopted an evolutionary approach by comparing the conservation of SRSF2 motifs between human and mouse *MDM2*. We evaluated conservation based on both sequence similarity and positional alignment. Our analysis revealed an intriguing pattern across exons, the sum of all conserved predicted SRSF2 binding-site score was elevated in terminal exons 4, 5, 10 and 11 but decreased in middle exons 6, 7, 8 and 9. Exons 4, 5, 10 and 11 displayed particularly high scores for conserved sites reinforcing the notion that terminal exons might preferentially be regulated by SRSF2 for damage-induced splicing (Supplementary Table 4). However the score for exon 10 was lower when taking exon length into account (0.045 compared to 0.083 and 0.088 for exons 4 and 11), perhaps explaining its lack of regulation in response to damage treatment in the minigene analysis.

### Point mutation at the mouse Srsf2 binding site in exon 11 causes exon exclusion and demonstrates conserved splicing regulation

To further examine the role of SRSF2-mediated regulation of the *MDM2* regulon, we turned to exon 11 for which we have previously identified both positive and negative splice regulatory elements (13, 24). To determine whether mutations in the Srsf2 binding site within exon 11 can regulate the splicing of preceding 6 exons (exon regulon), we performed mutational analysis of the Srsf2 binding site using the mouse *Mdm2* minigene . We previously showed that exon 11 in mouse *Mdm2* also contains Srsf2 binding sites that promote FL-*Mdm2* expression (24). We sought to mutate this Srsf2 binding site in mice to induce endogenous expression of *Mdm2-MS2*, the murine ortholog of human *MDM2-ALT1*. To tackle this, we designed Srsf2 mutations in the mouse *Mdm2* minigene that would decrease the predicted binding score of Srsf2 but would still maintain the same amino acid sequence encoded with exon 11. As discussed previously, we used the ESE finder tool (6) to determine the predicted Srsf2 binding motifs and their relative position on *Mdm2* Exon 11 (Figure 3A). Using this information, we designed silent mutations that would alter the predicted binding of Srsf2 to exon 11 but not affect the amino acid sequence of the resulting *Mdm2* protein. These mutants were generated by introducing single-nucleotide substitutions into the mouse *Mdm2* minigene using site-directed mutagenesis. Two of these mutations, C3A and C12G produced the largest decrease in the ESE predicted binding score dropping from 2.73 to 1.72 for C3A and 2.73 to 1.69 for C12G (Figure 3A (table)). Despite these reductions in predicted Srsf2 affinity, neither mutation noticeably altered exon 11 inclusion in the cells under normal or genotoxic stress inducing UV-C conditions (Figure 3B, C). Interestingly, a single nucleotide change at the seventh cysteine (C7A) in the first Srsf2 binding site of exon 11 caused a pronounced increase in exon 11 exclusion in the absence of stress which did not change a lot following UV treatment, despite having the smallest effect on the predicted binding score of Srsf2 (2.73 to 2.35) (Figure 3A, B, C). To further assess whether the C7A phenotype observed in our analysis was due to loss of Srsf2 binding at this site, we performed a rescue experiment using minigenes with additional nucleotide substitutions at the same Srsf2 predicted binding site (Figure 3D). In this case, we introduced an additional mutation at the sixth thymine (T6C) that increased the ESE score from 2.35 to 4.08. To assess the role of the T6C mutation independently of C7A, we also created a single T6C mutation, which raised the predicted Srsf2 binding score from 2.72 to 4.46. The double mutation, resulted in an almost complete rescue of the full-length isoform in both normal and UV conditions, whereas the single T6C mutation alone inhibited induction of the exon 11 excluded isoform, even in response to damage induction, supporting the role for Srsf2 binding in promoting exon retention (Figure 3E, F).

**Figure 3:**
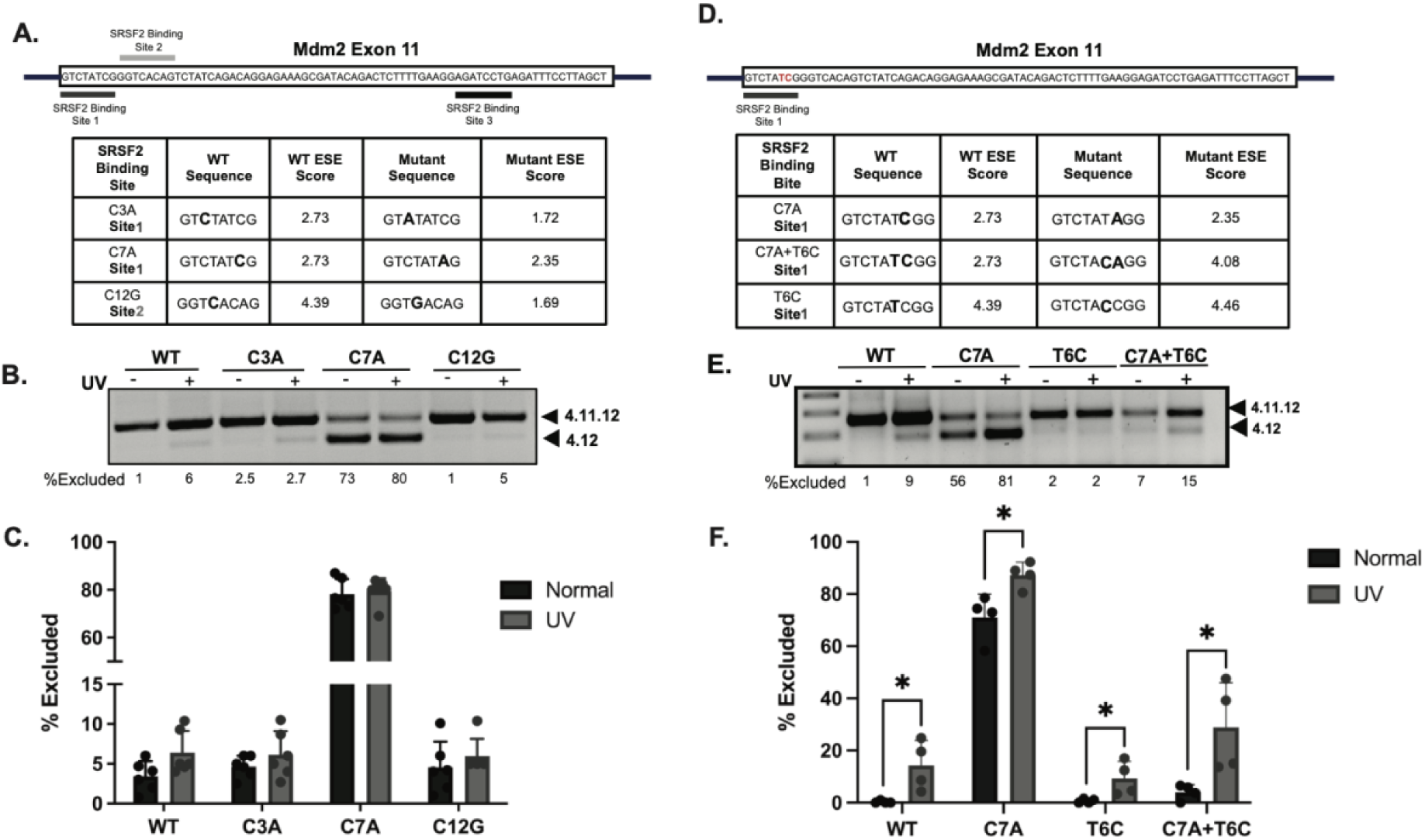
Point mutations at the Srsf2 binding site in exon 11 generates Mdm2-MS2 and demonstrates endogenous splicing regulation by this terminal exon. A. Schematic representation of *Mdm2* exon 11 showing predicted Srsf2 binding motifs (Sites 1, 2 and 3) and their relative positions using ESE finder. Table showing wild type (WT) and mutant exon splicing enhancer (ESE) sequences with calculated ESE scores for each Srsf2 binding site mutation. **B.** RT-PCR analysis of NIH3T3 cells transfected with WT mouse *Mdm2* 4-11-12 minigene as well as minigenes with respective mutations (C3A, C7A, C12G), treated with and without UV-irradiation. Exon 11 inclusion (4.11.12) and skipping (4.12) isoforms are indicated with percent exon exclusion (% excluded) being quantified below each lane. **C**. Quantification of n=3 independent experiments showing exon 11 skipping from fig. 3C under normal and UV-treated conditions. **D.** Summary of predicted ESE scores for WT and mutant Mdm2 constructs containing compensatory mutations (C7A, T6C, and C7A+T6C) at SRSF2 binding motifs. **E.** RT-PCR analysis of splicing products from mouse *Mdm2* exon 11 minigenes containing the indicated mutations, transfected into cells with or without UV-treatment. Percent exon exclusion is quantified below each lane. **F.** Quantification of exon 11 exclusion from minigene assays. The C7A mutation caused strong exon skipping under basal conditions, whereas T6C or C7A+T6C partially restored exon inclusion after UV-irradiation.

### A CRISPR-induced mutation at the Srsf2 binding-site within exon 11 generates Mdm2-MS2 and demonstrates endogenous splicing regulation

To assess whether this single mutation in exon 11 alone can generate the *Mdm2-MS2* isoform endogenously and more importantly, to assess its biological function, we decided to generate a CRISPR cell line with the silent C7A mutation. Initially we chose two different cell lines: one with an intact p53 pathway (C2C12) and one with an aberrantly functional p53 pathway (NIH3T3-ARF mutated). While we successfully generated the NIH3T3 cell line carrying the CRISPR-induced mutation, we were unable to isolate any C7A positive clones for the C2C12 cell line. RT-PCR analysis of our CRISPR-mutated NIH3T3 cells showed increased expression of *Mdm2-MS2* compared to control cells under both normal and UV-treated conditions. These results indicate that a single mutation at the Srsf2 binding site within exon 11 is sufficient to alter canonical *Mdm2* splicing and promote endogenous production of *Mdm2-MS2* (Figure 4A, B). Additionally, since we planned to induce this mutation using a homology directed repair (HDR) template in mice, it was necessary to incorporate a second synonymous mutation within *MDM2* to disrupt the PAM (protospacer adjacent motif) site to prevent repeated Cas9 cutting and improve the efficiency of gene editing. To determine the relative effect the PAM mutation would have on the splicing of exon 11, we analyzed the binding matrices using ESE finder. We found that changing the guanine to adenine at position 9 (G9A) of exon 11 within the PAM site, does not affect the binding score of the first Srsf2 binding site, but decreased the binding score of the second Srsf2 binding site from 4.39 to 2.64 (Supplementary Figure 1A). To experimentally test the functional consequence of the additional mutation, we generated a new mouse minigene carrying both mutations (C7A; G9A) and transfected them into NIH3T3 cells. After 24-hours, we treated these cells with UV-C to induce genotoxic stress and to assess the expression of the skipped isoform using RT-PCR. Our data shows that the splicing effect on exon 11 is essentially identical between the single C7A mutation and the double C7A; G9A mutation, both mutations causing a significant reduction in exon 11 inclusion in the absence of stress (Supplementary Figure 1B).

**Figure 4:**
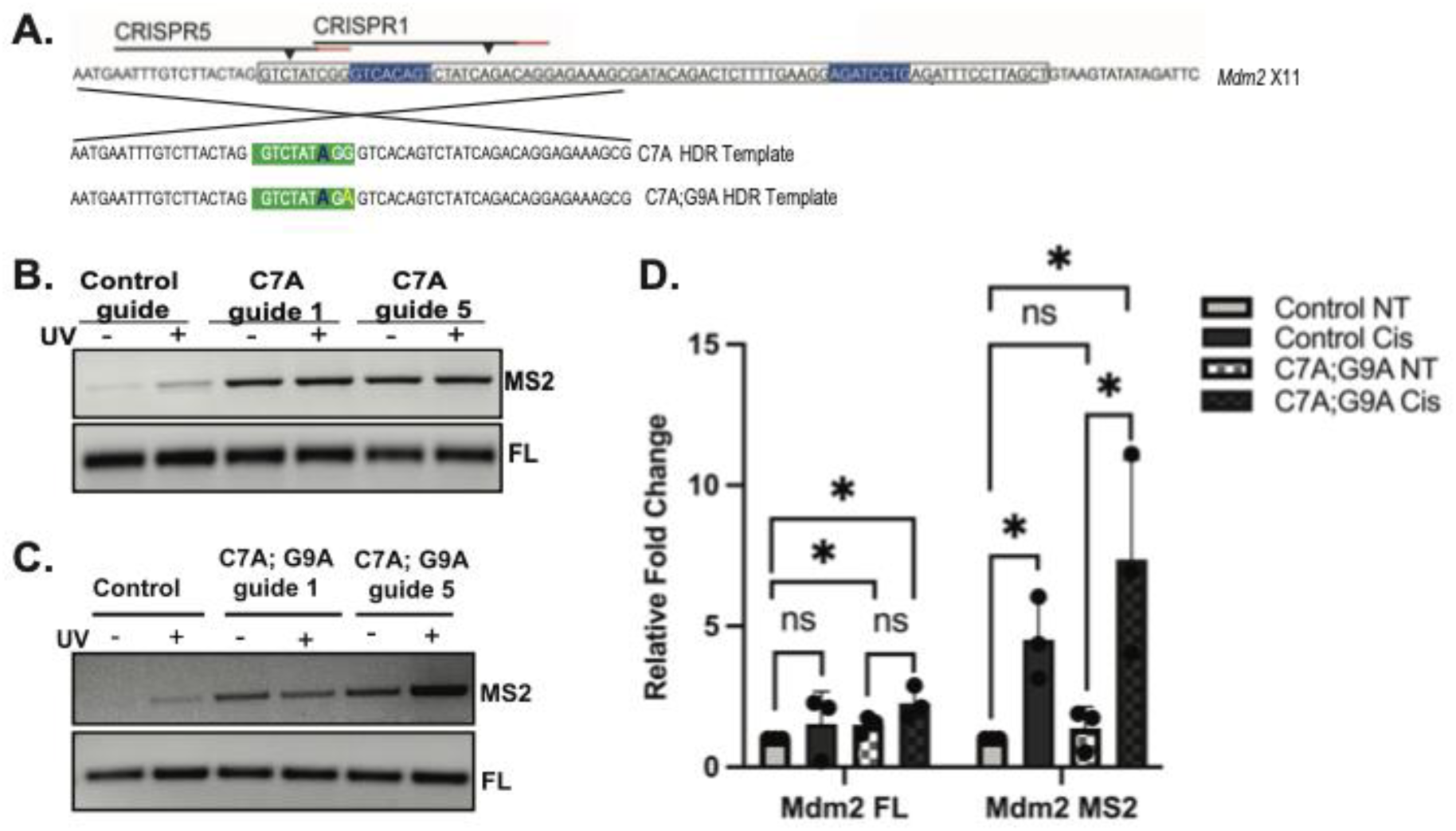
A CRISPR-induced mutation at the Srsf2 binding-site within exon-11 generates *Mdm2-MS2*. **A.** Schematic representation of guide RNA (gRNA) and homology-directed repair (HDR) template design used for introducing the C7A;G9A mutation in *Mdm2* exon 11 of NIH3T3 cells. **B, C.** RT-PCR analysis showing endogenous expression of *Mdm2-MS2* and *Mdm2-FL* transcripts in control, C7A alone and C7A;G9A double mutant NIH3T3 cells with and without UV-treatment. **D.** qPCR analysis of *Mdm2-FL, Mdm2-MS2* and *Mdm2* total splicing in control and C7A;G9A double mutant NIH3T3 cells with and without cisplatin-treatment.

We used two CRISPR guides to generate CRISPR-mutated NIH3T3 cell lines harboring both the C7A and G9A mutations (Figure 4A). To assess the effect of the double point mutations and the induction of *MS2*, cells were seeded and treated with genotoxic stress for 24 hours. Genotoxic stress was induced using either treatment with UV-irradiation or cisplatin as both stressors cause DNA helix distorting damage and promote the expression of *Mdm2-MS2* in mice or *MDM2-*ALT1 in humans. After 24 hours post-treatment, RNA was harvested from these cells to assess *Mdm2* splicing patterns. RT-PCR analysis revealed that our new CRISPR-modified cell lines expressed *Mdm2-MS2* even in the absence of genotoxic stress (Figure 4C). qPCR data further corroborated this showing that in the presence of the C7A;G9A mutation, *Mdm2-MS2* levels are higher compared to the wildtype control cells and are further increased after cisplatin treatment (Figure 4D). These data demonstrate that the splicing effects observed in the mutated minigene system are recapitulated endogenously. Additionally, NIH3T3 cell harboring CRISPR-mediated C7A and G9A mutations endogenously expressed *Mdm2-MS2,* supporting the feasibility of generating a C7A;G9A CRISPR mouse model.

### Constitutive endogenous expression of Mdm2-MS2 increases proliferation and decreases apoptosis in cells lacking an intact p53 pathway

To analyze the functional consequences of induced *Mdm2-MS2* expression in cells, we subjected control and C7A; G9A CRISPR mutated NIH3T3 cells to proliferation assays. Briefly, we seeded equal number of control and C7A; G9A mutant cells and placed them in the IncuCyte live-cell imaging system to measure cell-density to infer proliferation rates. We predict that introducing *Mdm2-MS2* would decrease proliferation of wildtype p53 cells but increase proliferation in cells where p53-pathway is inactivated. The C7A;G9A CRISPR-mutated NIH3T3 cells showed significantly increased cell confluency compared to the control cells, indicating a higher proliferation rate (Figure 5A). We next examined whether the mutation also affected apoptosis rate. To measure apoptosis, we treated the cells with cisplatin and measured the number of Caspase3/7 positive cells marked by a green-fluorescent dye that intercalates with the DNA of apoptotic cells. After a 72-hour time period, we found that the C7A;G9A CRISPR mutated cells had decreased apoptosis in response to cisplatin treatment indicating a resistance to apoptosis (Figure 5B). Furthermore, to assess the transformation potential of these cells, we performed a colony formation assay. We found that while the control cells formed an average of 5.33 colonies per 2 cm dish, C7A;G9A mutated cells were able to form 23.67 colonies on average (difference in mean 18.33; SEM ± 5.24) (Figure 5C), indicating an increase in clonogenic capacity. Given the observed changes in proliferation, apoptosis, and transformation of these cells, we were curious about the downstream genes affected by *Mdm2-MS2* expression, therefore we performed qRT-PCR to analyze the expression of p53 transcriptional targets in control and C7A;G9A CRISPR mutated cells (Figure 5D). Of these targets, we found that *CCND1 (Cyclin D1),* a gene involved in cellular growth, metabolism and differentiation to be significantly upregulated in the *Mdm2-MS2* overexpressing cells (Figure 5D) (31,32). The expression of other canonical p53 target *P21* is also increased however targets such as *Puma*, *Bax* and *Killer* were not changed in these C7A; G9A CRISPR mutated cells. Our data shows that the C7A;G9A mutation in NIH3T3 cells increases the *Mdm2-MS2* expression and imparts a proliferative phenotype to P19-ARF mutant cells. These findings contrast with the expectations we had for wildtype cells with an intact p53 pathway. In wildtype cells we hypothesize that increased expression of *Mdm2-MS2* would lead to cell cycle arrest or apoptosis, by alleviating the *MDM2* mediated repression of the p53 tumor suppressor. These data suggest that the integrity of the p53-pathway plays a critical role in determining the functional impact of *Mdm2-MS2* expression.

**Figure 5:**
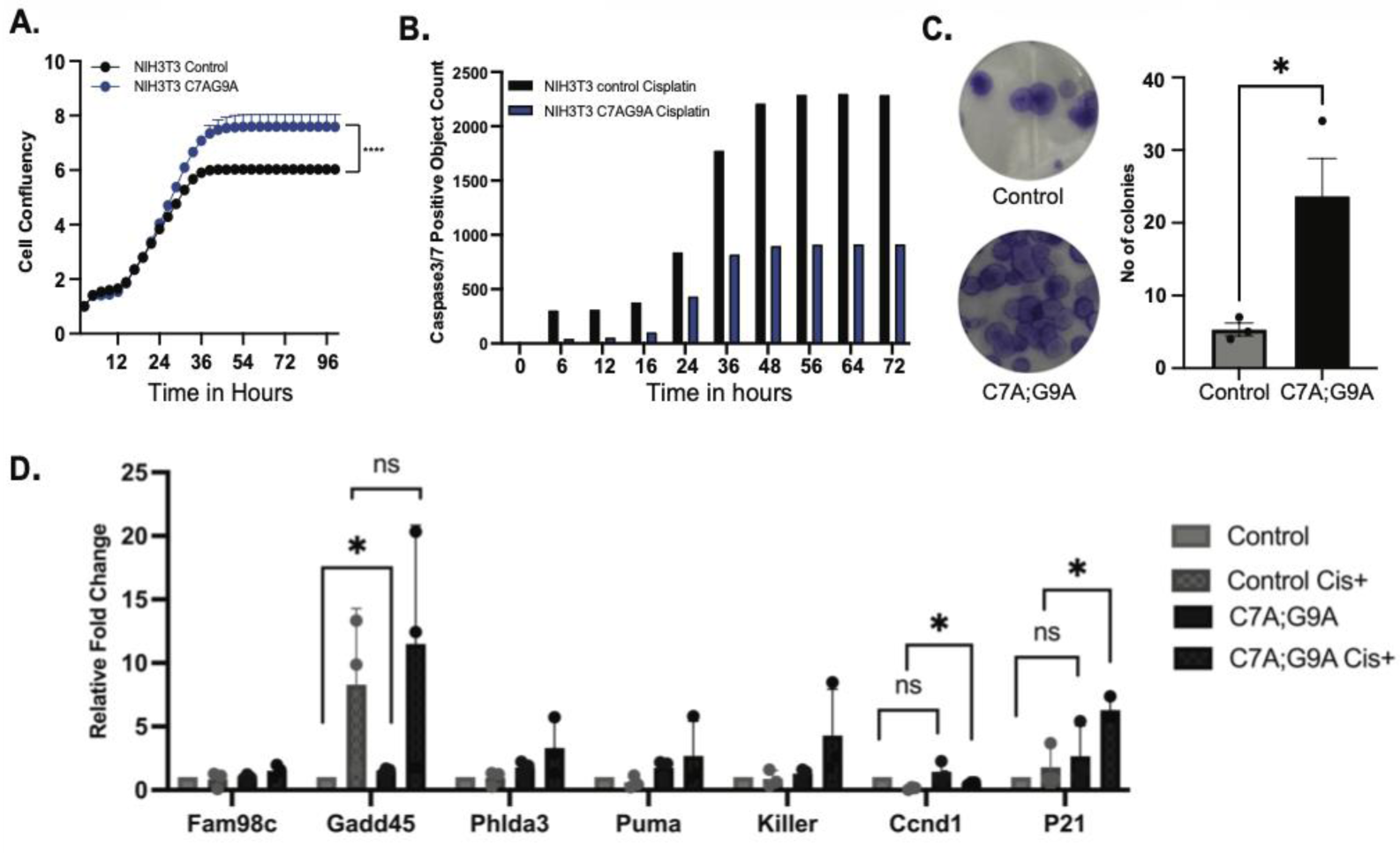
The constitutive endogenous expression of *Mdm2-MS2* increases proliferation and decreases apoptosis in cells without an intact p53 pathway. A. NIH3T3 control and C7A;G9A mutant cells were subjected to proliferation in the IncuCyte, and growth was measured over a period of 5 days. **B**. NIH3T3 control and C7A;G9A mutant cells were treated with 75uM cisplatin and IncuCyte ^®^ Caspase3/7 dye was added. When added to tissue culture medium, the inert, non-fluorescent substrate crosses the cell membrane where it is cleaved by activated caspase 3/7, resulting in the release of the DNA dye and fluorescent staining of the nuclear DNA and the flourorescense positive cells are measured by the incucyte. **C**. Colony formation assay shows that the C7A;G9A mutant cells can form more colonies than the wild type cells. **D**. RT-PCR shows changes in the expression of some p53 target genes in C7A;G9A mutant cells.

### Utilizing Crispr to generate a novel mouse model that overexpresses the splice isoform Mdm2-MS2

Previously, our lab has reported a transgenic mouse model that expresses the human *MDM2-ALT1* isoform from a transgene (33). In these p53-null mice, the expression of *MDM2-ALT1* (the human isoform of *Mdm2-MS2*) caused increased tumorigenesis and shifted the tumor spectrum towards rhabdomyosarcoma (33). This *MDM2-ALT1* transgenic model artificially overexpresses the human MDM2-ALT1 protein but, does not mimic the actual splicing change in endogenous *Mdm2* pre-mRNA, nor does it avail an opportunity for splice altering therapies. To circumvent this, we decided to take advantage of our findings thus far and mutate the Srsf2 binding-site in *Mdm2* exon 11 using *CRISPR*-*Cas9* gene editing to generate a p53-wild type mouse model with altered *Mdm2* splicing. In our endogenously induced splicing model, we hypothesize that the *Mdm2 mice* will give rise to the *Mdm2-MS2* isoform as a result of the mutated Srsf2 binding-site in exon 11. To test our hypothesis, we employed the same *CRISPR-Cas9* genome engineering strategy previously described (Figure 4) and induced the C7A;G9A point mutations into mouse embryos to generate five founder mouse lines of a novel constitutively *Mdm2*-*MS2* expressing mouse model. Taking advantage of the unique enzymatic SfcI restriction site C7A;G9A generates, we employed a RFLP-PCR (Restriction fragment length polymorphism) as the primary genotyping strategy. Following PCR amplification, we are able to digest the product using SfcI which only cleaves mouse DNA harboring the C7A;G9A mutations (Supplementary Figure 2A, B). Of the five founder mouse lines, were able to breed three founders to generate pups. After RFLP-PCR genotyping of the mice, we amplified the *Mdm2* region carrying the suspected mutation and conducted Sanger sequencing as a secondary method of verification of the mutation (Supplementary Figure 2C). After breeding all three founder mouse lines we were able to confirm the mice were born with accurate Mendelian ratios indicating that there was no resultant embryonic lethality phenotype (Supplementary Figure 2D).

### Mdm2 exon 11 C7A; G9A Mice Show Increased Mdm2-MS2 Isoform expression and increased expression of P53 target genes

After we confirmed that the mice carried the C7A;G9A mutation in *Mdm2* exon 11 (Supplementary Figure 2), we examined the expression of the *Mdm2-MS2* isoform in mouse embryonic fibroblasts (MEFs) from all three founder lines. To assess the *MS2* levels in mutant MEFs and to determine whether they are damage responsive, we treated them with UV-irradiation and performed RT-PCR on the RNA harvested from these MEFs. We found that mutant MEFs from all three lines had an increased expression of *Mdm2-MS2* under normal conditions that was further increased after UV-irradiation (Figure 6A). This increase in *Mdm2-MS2* after UV-irradiation was minimal in lines 3 and 14 but was well demonstrated in line 5 MEFs.

**Figure 6:**
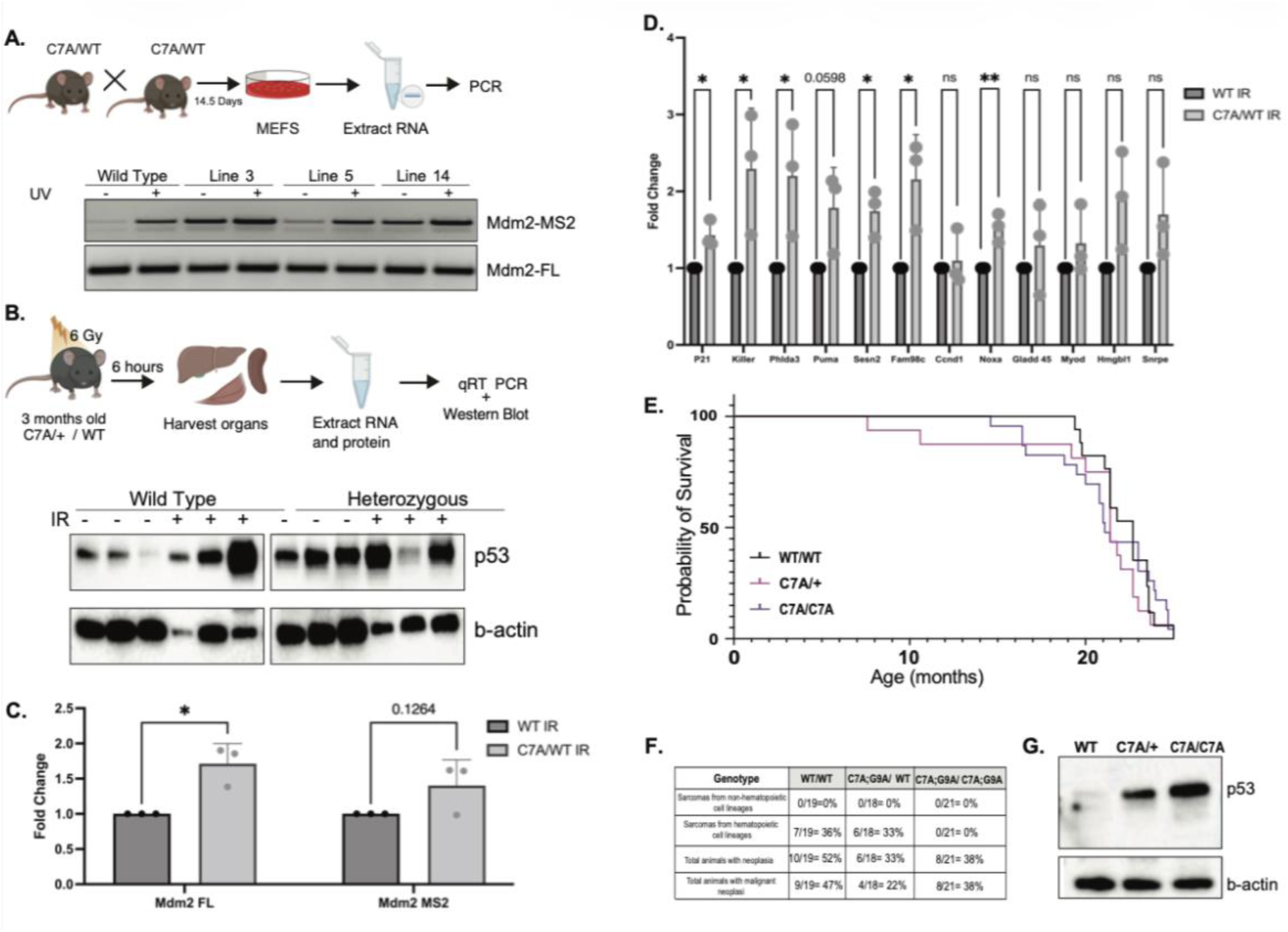
*Mdm2* exon 11 C7A; G9A mice show increased *Mdm2-MS2* isoform expression and increased expression of p53 target genes. **A.** RT-PCR of Mouse embryonic fibroblasts (MEFs) for *Mdm2-FL* and *Mdm2-MS2* in control, and UV-C treated cells. **B**. Schematic showing mice irradiation proceedure, line 5 mice were treated with 6 Gy gamma irradiation and harvested after 6-hours followed by protein and RNA isolation. Western blot was preformed to analysize p53 protein levels, showing increased levels in of p53 in C7A;G9A heterozygous compared to wild type and increased p53 levels for both wild type and C7A;G9A heterozygous after gamma irradiation. **C.** RT-PCR for mouse *Mdm2-FL* and *MS2* in spleen harvested from mice treated with gamma irradiation. **D**. qPCR for *p53* target genes on spleen of wild type and C7A;G9A gamma irradiated mice. **E, F**. Kaplan Meier curves and tabular representation of wild type, C7A;G9A heterozygous and C7A;G9A homozygous mice numbers showing incidence of sarcomas and hematopoietic sarcomas in these mice. **G**. Western blot analysis showing p53 protein expression in wild type, heterozygous and homozygous mice.

Given that *MDM2-ALT1* can bind full-length *MDM2* and stabilize p53 by preventing the *MDM2* mediated ubiquitination of p53, we next investigated whether our mouse model has any altered p53 response. We irradiated both line 5 *Mdm2 exon 11* C7A;G9A mice, as well as sibling control mice with 6-Gy gamma irradiation (34) and harvested spleen which is highly responsive to radiation exposure for protein and RNA 6-hours post-treatment (Figure 6B). Though p53 protein expression is noticibly higher in the C7A;G9A mice in the absence of any treatment, we see a clear increase in p53 levels in 2 of the 3 mutant mice after irradiation (Figure 6B). qPCR analysis from irradiated mice demonstrated that the C7A; G9A mutant mice had elevated *Mdm2-FL* and a marginally increased *Mdm2-MS2* expression compared to the wild-type mice (Figure 6C). Given this mouse model was generated in a p53-wild type background, we next examined p53 target expression using qPCR to determine whether altered *Mdm2* splicing changed p53’s ability to regulate its target genes. Our data shows that in a p53 WT setting, irradiation in C7A; G9A mutant mice causes an upregulation of p53 target genes, including apoptosis targets *puma*, *noxa* and *killer* among others (Figure 6D). To account for potential loss of signal at 6 hours, we also performed a 3-hour post 6-Gy gamma irradiation tissue harvest in our line 3 *Mdm2 exon 11* C7A;G9A mice (Supplementary Figure 3). With the exception of *p21*, this upregulation is exagerrated compared to the wild type induction of these same targets indicating that there is an enhanced p53 response in *MS2* expressing mice.

To investigate the long-term consequences of *Mdm2-MS2* overexpression in an *in vivo* setting, we generated a cohort of wild type and *Mdm2 exon 11* C7A;G9A mice and monitored them over 24-months to determine if the constitutive expression of *Mdm2-MS2* leads to any phenotypic differences. Given the intact p53-pathway in these mice, we anticipated that sustained expression of *Mdm2-MS2* might confer protection against tumor development, since *Mdm2-MS2* should stabilize p53 levels. However, no difference in lifespan were observed and the tumor onsets were comparable across genotypes (Figure 6E). On the other hand, comparative pathological evaluation determined that, in the absence of an external insult, none of the mice developed any sarcomas originating from non-hematopoietic cell lineage. This lack of background tumorigenesis provides a controlled context in which we can assess the impact of altered *Mdm2* splicing on hemopoietic transformations. Strikingly, *Mdm2 exon 11* C7A;G9A homozygous mice developed significantly fewer sarcomas from hematopoietic origin (0/21=0%) compared to heterozygous (6/18=33%), and wild type mice (7/19=36%) suggesting that *Mdm2-MS2* overexpression is protective against these types of malignancies (*P-value* wild type vs C7A;G9A homozygous=0.0081). Additionally, we observed that all three groups of mice developed age-related neoplasia, but specifically 47% of wild type mice developed malignant neoplasia compared to only 22% of C7A;G9A heterozygous and 38% of C7A;G9A homozygous mice, again pointing to the protective role of *Mdm2-MS2* overexpression in the p53 wild type background (Figure 6F). In summary, our data demonstrates that mutation of the Srsf2 binding-site in exon 11 in an *in vivo* system leads to the constitutive expression of *Mdm2-MS2*, independent of genotoxic stress and that constitutive expression leads to increased levels of p53 protein (Figure 6D, G). Furthermore, these mice harboring the C7A;G9A genotype, demonstrate partial protection against age-induced neoplasia, highlighting the functional significance of regulated splicing in tumor suppression.

## Discussion

Alternative splicing of *MDM2* plays an important role in regulating p53 signaling, stress response and cancer development, yet the mechanisms governing this process are not well understood. This manuscript focuses on understanding a part of this mechanism by asking two key questions: 1. is the multi-exon cassette of *MDM2* excised as indiuidual exons or as a coordinated eight-exon unit and 2. what are the functional consequences of constitutive *MDM2-ALT1* expression? In silico analyses of splice-site strength and SRSF2 binding motifs, together with a comprehensive modular minigene system designed to interrogate splicing of individual *MDM2* exons under genotoxic stress, demonstrate primarily that intronic context is essential for exon recognition, and once recognized, only exons 4, 5, and 11 are responsive to DNA damage. These terminal exons exhibit higher predicted SRSF2 binding-site scores compared with exons 6–9, suggesting that exons 4, 5, and 11 act as focal points for SRSF2-dependent splicing regulation. Collectively, these findings support an “exon regulon” model for *MDM2* splicing under genotoxic stress, in which terminal exons are subject to heightened regulation by both cis- and trans-acting elements, resulting in coordinated excision of the eight-exon cassette rather than independent exon skipping (Figure 7). It should be noted that alternative splicing of *MDM2* gives rise to multiple spliced isoforms only one of which is interrogated extensively in this paper.

**Figure 7:**
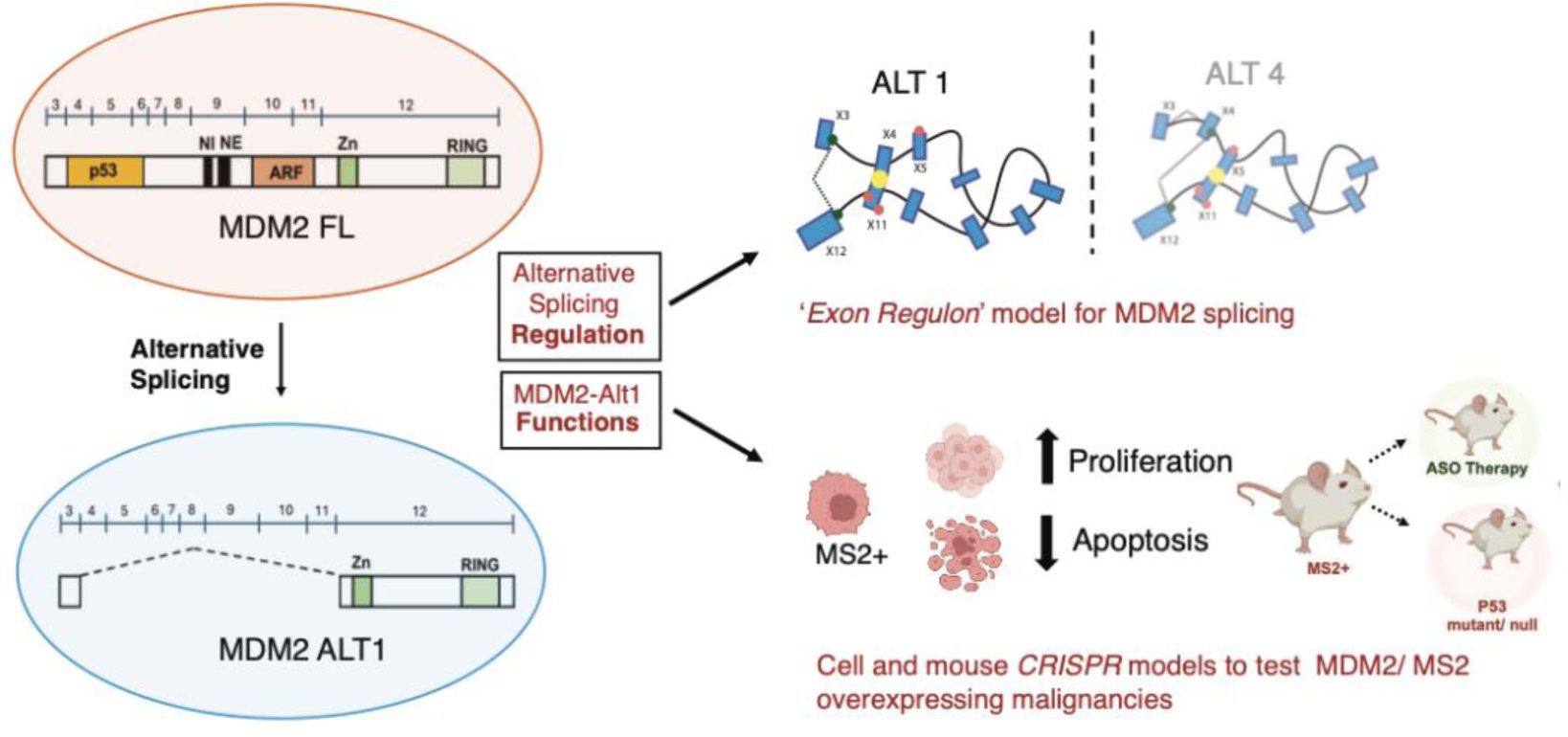
Graphical model representing the splicing of *MDM2*. Schematic depicting *MDM2* alternative splicing and how the regulon model could lead to distinct *MDM2* spliced isoforms. *MDM2-ALT1* overexpressing *CRISPR*-engineered cells and mice could serve as models delineating the biological functions of *MDM2-ALT1/Mdm2-MS2* in vivo.

As the field of alternative splicing has expanded substantially over the past decade, these findings are of particular interest for several reasons. i) Increasing evidence suggests that splicing decisions are not made independently at each exon, but instead can involve coordinated processing of exon blocks through higher-order RNA organization and long-range interactions. Structural rearrangements of the pre-mRNA, driven by RNA secondary structure or protein-mediated looping, may bring non-adjacent cis-elements into proximity, thereby enabling coordinated inclusion or exclusion of multi-exon cassettes. Recent work of Bergfort et al. has highlighted the concept of exon blocks, in which groups of exons behave as modular regulatory units rather than as isolated entities, providing a framework for understanding how large exon cassettes can be excised in a concerted manner (35–37). In this context, regulated exclusion of *MDM2* exons 4–11 may reflect an ordered or hierarchical splicing process, potentially involving recursive splicing events that are selectively engaged under stress conditions (38,39). Such mechanisms would allow distal terminal exons, such as exons 4 and 11, to act as regulatory anchors that dictate downstream splicing outcomes, consistent with our exon regulon model and recent emerging paradigms in splicing regulation.

ii) As the splicing field increasingly focuses on how regulatory networks govern exon inclusion and exclusion, our findings support the emerging view that specific exons can function as focal points for splicing regulation. Glidden *et al.* demonstrated that exonic variants capable of disrupting splicing are non-uniformly distributed across the genome and they identified a subset of exons termed “hotspot exons”, that are particularly susceptible to splicing perturbations, including those induced by pharmacologic treatments (40). We interrogated this ‘hot-spot’ exon data set and found that *MDM2* exon-11 is not classified among the “hotspot exons” identified in large-scale analyses of splicing perturbations, indicating that this exon (exon 11) is not broadly susceptible to generic splicing disruption but is instead regulated in a highly context-dependent manner. This distinction underscores that stress-induced splicing of *MDM2* is not driven by intrinsic exon fragility, but by dedicated cis-regulatory elements and stress-responsive trans-acting factors that selectively reprogram splicing decisions. Our work therefore highlights that alterations in cis-regulatory elements can impact splicing outcomes, suggesting that exon-specific therapeutic strategies, such as antisense oligonucleotides, may provide greater specificity than global splicing modulation.

iii) An additional layer of complexity in regulated *MDM2* splicing may arise from the order in which splicing events occur and the influence of distal regulatory elements on exon definition. In addition to cis-regulatory elements and splicing factors, the order and interdependence of intron removal can play a critical role in shaping alternative splicing outcomes. Genome-wide analyses and nascent RNA profiling reports have demonstrated that introns are often removed in a defined, transcript-specific order, and that splicing of one intron can influence the processing of neighboring introns, particularly in transcripts undergoing regulated exon skipping (41–43). Such intra-dependencies provide a mechanistic framework for coordinated multi-exon exclusion events like that of *MDM2*, and support our findings that *MDM2* exon skipping is not stochastic but instead reflects ordered and structured splicing decisions involving distal regulatory elements.

iv) Additionally and within this framework, ‘exon definition’ provides a compelling mechanism for how SRSF2 may exert control over coordinated *MDM2* splicing decisions. Exon definition relies on early recognition of exon boundaries through cooperative interactions between splice-sites and exon-bound splicing factors, a process that is particularly important in complex mammalian genes (44,45). Consistent with this model, SRSF2 engages conserved ESEs within *MDM2* exon 11 and promotes its inclusion under basal conditions, indicating that SRSF2 helps to define exon 11 as a splicing unit. Disruption of SRSF2 binding, either through genotoxic stress or cis-element mutation, would be expected to weaken exon 11 definition, favoring an alternative pairing of splice-sites that yields coordinated skipping of the exon 11–containing cassette and production of *MDM2-ALT1*. This provides a mechanistic link between exon definition, stress-responsive modulation of SR-protein activity, and regulated exclusion of multiple *MDM2* exons.

Regardless of the mechanism, the fact that an array of *MDM2* isoforms arise in response to different types of stress and impart distinct changes in the p53-pathway suggests that splicing control of *MDM2* acts as a rheostat to control p53 response. Given the central role of the p53 pathway in cancer biology, and its inhibition by overexpression of *MDM2*, manipulation of *MDM2* splicing represents a potential strategy for reactivating p53’s tumor-suppressive activity and holds significant therapeutic promise. With this therapeutic potential in mind, our lab investigated the functional consequences of endogenous *MDM2-ALT1* expression, by generating a CRISPR-modified NIH3T3 cell line with silent mutations in the conserved Srsf2 binding-site of exon 11. In our engineered system, mutations in the exon 11 cis-elements were sufficient to promote skipping of the internal exon cassette and generate the *Mdm2-MS2* isoform, but this occurred without a detectable reduction in the endogenous *Mdm2-FL* transcript. This differs from the DNA damage response, where genotoxic stress activates p53, increases *Mdm2* transcription, and concomitantly redirects splicing of the endogenous pre-mRNA toward *Mdm2-MS2*, leading to a relative loss of *Mdm2-FL*. In contrast, constitutive expression of *Mdm2-MS2* from an exogenous construct generates the alternative isoform independently of the native promoter and without globally reprogramming splicing at the locus, so it does not compete with or diminish production of the full-length transcript.

Functionally, our *Mdm2-MS2* CRISPR mutant cells demonstrated dysregulated proliferation and apoptosis as well as changes in multiple p53 target genes. Of note, *CCND1* is upregulated and its known function as a driver of cell cycle progression correlates well with the increased proliferation phenotype seen in NIH3T3 cells. To extend these findings *in vivo*, we generated a CRISPR mouse model that expressed *Mdm2-MS2* across all tissues under homeostasis. These *MDM2-exon 11* CRISPR mice demonstrated a protection against age-induced neoplasia suggesting that constitutive *in vivo* expression of *Mdm2-MS2* may be protective in a p53 wild-type background. In *Mdm2-MS2* CRISPR mutant cells, p53 target genes were not upregulated however in our CRISPR mutated mice where the p53-pathway is intact, we observed a significant upregulation of p53 target genes, exceeding that in the wildtype mice, upon gamma irradiation. Together, these finding highlight our *Mdm2* exon 11 C7A; G9A mouse model as powerful tool for studying the long-term consequences of splicing dysregulation *in vivo* both within a functional p53 context, as done in this work, and in the presence of p53 mutations by crossing our mice to distinct p53 mutant-lines.

The biological relevance of splicing regulation in cancer is underscored by extensive evidence that aberrant alternative splicing contributes to tumorigenesis and progression across multiple cancer types. Dysregulated splicing events can generate isoforms that promote proliferation and survival, or that impair apoptotic responses, thereby functioning as drivers of oncogenic pathways rather than mere byproducts of malignant transformation. Aberrant splicing has been linked to alterations in tumor suppressors and oncogenes, and disruptions in splicing regulatory factors themselves are observed in tumors, emphasizing that splicing perturbations can directly affect cancer hallmarks such as cell cycle control and apoptotic resistance (2,46–49). These cancer-associated splicing changes often arise from both mutations within splicing regulatory elements and altered expression or activity of splicing factors, reinforcing the concept that precise control of splice-site choice is critical for maintaining normal cellular homeostasis (50,51). Clinically, when a SNP is identified in an exon or intron, splicing effects are typically evaluated only for the immediately adjacent exons; our work suggests that such localized assays should be complemented by analyses of splicing across the full transcript to uncover broader splicing defects that would be missed by a narrowly focused approach.

By situating the *Mdm2* exon 11 model within this broader landscape, we show that regulated splicing of a key proto-oncogene is not simply a molecular curiosity but reflects a mechanism by which cells utilize alternative splicing mechanisms to fine-tune stress responses with direct consequences for tumor biology. These insights further support the utility of splicing models to dissect how *specific* splicing events shape cancer-relevant pathways and highlight alternative splicing as a potential avenue for targeted modulation in diseases such as cancer (52).

## Materials and Methods

### Construction of *MDM2* minigenes

All minigene constructs were derived from a previously constructed 3-11-12 minigene (Singh, 2009) cloned into the pCMV-tag 2B vector. For the minigene system shown in Figure 1, exon 3 was PCR-amplified using primers containing overhangs corresponding to the 3’ splice-site of the intron adjacent to the intermediate exons (exons 4 through10). Similarly, exon 12 was amplified with specific primers. The intermediate exons (4 to 10) with introns containing 10 nt native intronic sequences on the 5’ and 3’ end, were amplified using primers with reverse overhangs complementary to the 5’ end of the forward primer used for exon 12 amplification. This strategy resulted in PCR-amplified fragments with complementary overhangs. Subsequently, the three fragments were joined through a three-piece PCR ligation reaction. The resulting construct was then cloned into the pCMV-tag 2B vector and sequenced using Sanger sequencing (Figure 1A). Figure 2 minigenes were designed with custom dsDNA inserts purchased from Integrated DNA Technologies (gBlocks Gene Fragments). These inserts contained an *MDM2* exon (4 to 10) with ≥ 90 bp of the native 3’ end intronic sequence upstream of the exon and ≥ 35 bp of the native 5’ end intronic sequence downstream of the exon (See Figure 2, for specific intron lengths). All inserts contained the necessary spliceosome recognition sequences to facilitate biologically accurate splicing behavior; *MDM2* sequences were referenced from Ensembl (ENST00000258149.11). Each insert was cloned into the 3-11-12 *MDM2* minigene construct by linearizing the 3-11-12 minigene at the EcoRI and PstI restriction-sites and adding the insert using the in-fusion cloning protocol with primers designed to include a 15 bp overlap upstream of the PstI-site and downstream EcoRI-site of the linearized construct (see table of primers). Final constructs retained both restriction sites, EcoRI and PstI and their sequences were confirmed via Sanger sequencing.

### Transient transfection and splicing evaluation of minigenes

500,000 HeLa cells were seeded in 6-well plates. After 24 hours, 1 µg of minigene construct was transiently transfected using Lipofectamine 3000 (ThermoFisher catalog#: L300015). Cells were treated with either 50 J/m^2^ UV-C or 75 µM cisplatin 24 hours post-transfection and were harvested for RNA using the RNeasy Kit (Qiagen catalog#: 74106) 48 hours post-transfection. Reverse transcription PCR was run using a vector-specific FLAG-tag forward primer and an *MDM2* exon 12 reverse primer (Primers are listed in Supplementary Table 5).

### CRISPR-Cas9 genome editing

NIH3T3 and C2C12 cells were transfected with either a control plasmid SpCas9-2A-EGFP or SpCas9-2A-EGFP-*Mdm2* along with a HDR custom single-stranded DNA 243 base pair repair template. Four hours post-transfection, cells were treated with 1 mM SCR7 (Xcessbio). After 48 hours, cells were sorted for GFP expression on a BD Influx FACS cell sorter running SortWare software. GFP-positive cells were plated and maintained in a medium containing 1 mM SCR7. After two passages, NIH3T3 cells were collected for genomic DNA and genomic changes were verified by Sanger sequencing. We did not obtain positive clones for C2C12 cells.

### Generation of the mouse model

The *Mdm2 X11* C7A mouse model was generated by the Genetically Engineered Mouse Modeling Core of the Ohio State University Comprehensive Cancer Center. The synthetic single strand oligo donor DNA (ssODN) containing the C7A mutation together with the G9A PAM-site mutation was purchased from Integrated DNA Technologies (Coralville, Iowa, USA). Synthetic tracrRNA and crRNA was purchased from Sigma-Aldrich (Saint Louis, MO, USA). GeneArt Platinum Cas9 nuclease purified protein (cat# B25642) was purchased from Invitrogen (ThermoFisher Scientific; Waltham, MA, USA). The mix of assembled sgRNA with Cas9 protein (RNP complex) and ssODN was microinjected into C57Bl/6Tac zygotes. WT (wildtype) C57Bl/6Tac mice were used for founder mating purposes.

### Animal studies

All animal experiments were performed according to institute standards approved by the Institute Animal Care and Use Committee (IACUC) at The Abigail Wexner Research Institute at Nationwide Children’s Hospital and the Ohio State University. The *Mdm2 X11* C7A;G9A mouse model was generated by CRISPR/Cas9 technology as described above. Genotyping was performed by RFLP using *SfcI* (New England Biolabs, cat# R0561L) and *Mdm2* specific primers or by direct sequencing of PCR products. For P53 induction experiments, three 12-week-old *C7A* and three WT C57Bl/6 mice were irradiated with 6 Gy in a X-Rad 320 irradiator (Precision X-Ray, North Branford CT) and euthanized 3 or 6 hours later. Spleen tissue was collected and used to extract protein and RNA. Total protein extraction was performed using an Ultra-Turrax tissue homogenizer (IKA, Wilmington NC) in RIPA buffer, while total RNA extraction was performed using TRIzol reagent (Invitrogen, cat# 15596018) following the manufacturer’s instructions. Mouse embryonic fibroblasts (MEFs) isolation was carried out by harvesting 14.5-dayembryos from a *C7AxC7A* heterozygous crosses, in the presence of trypsin. Isolated MEF lines were maintained in DMEM medium and genotyped by RFLP or sequencing as described above.

Mice were monitored over a period of 24 months. Mice showing signs of pain or distress or with masses, abscesses and tumors more than 2.0 centimeters were humanely euthanized according to endpoint criteria. At 24-months, the mice from the *MDM2 X11* line 5 cohort were euthanized, necropsied, and examined for lesions and tissue abnormalities. Tissue specimens were fixed in 10% neutral buffered formalin, paraffin embedded, and sectioned, followed by hematoxylin and eosin staining and pathological analysis.

### RT and PCRs

Reverse transcription (RT) reactions were carried out using Transcriptor reverse transcriptase enzyme (Roche Diagnostics) with 1 µg of total RNA. Non-quantitative endogenous *MDM2* PCRs were performed as reported previously (22). *MDM2* minigene PCRs were performed as reported previously (23). *Mdm2* amplicons and PCRs for *MDM2* after RNA immunoprecipitation of T7-SRSF2 were performed using a set of nested primers under standard PCR conditions using Taq polymerase (Sigma Aldrich).

### Quantitative PCR

All Quantitative PCRs (qPCR) were performed with standard PCR conditions using a CFX384 Touch Real-Time PCR Detection System (BioRad). Real-time PCR reactions were carried out using the PowerUp SYBR Green PCR Master Mix (Applied Biosystems). The primers used to amplify the p53-target transcripts and *Mdm2-MS2* have been described in S1 table (Supplementary Table 1). All PCR reactions were carried out in triplicate experiments with three technical replicates for each and the amplification of single PCR products in each reaction was confirmed using a dissociation curve.

### Live cellular growth assay

Growth curves were generated from triplicate plating of either NIH3T3 Control CRISPR, NIH3T3 C7A;G9A CRISPR5 cell lines. Cells were seeded at a density of 5× 10^4^ cells per well in 12-well plates and analyzed for confluency using the IncuCyte live cell imaging system. Statistical significance was calculated by two-way ANOVA.

### Live cellular apoptosis assay

NIH3T3 control and CRISPR mutant cells were seeded at 10000 cells per well in 96-well plates. After 24-hours, cells were treated with cisplatin and placed in the IncuCyte to measure the number of Caspase3/7 positive cells marked by a green-fluorescent dye that intercalates with the DNA of apoptotic cells. Incucyte^®^ Caspase-3/7 Dyes for Apoptosis couple the activated caspase-3/7 recognition motif (DEVD) to a DNA intercalating dye. When added to tissue culture medium, the inert, non-fluorescent substrate crosses the cell membrane where it is cleaved by activated caspase-3/7, resulting in the release of the DNA dye and fluorescent staining of the nuclear DNA. With Incucyte^®^ integrated analysis software, fluorescent objects can be quantified and background fluorescence minimized.

### In-silico analysis: Supplementary Table 1

Table 1 was generated using data from multiple sources. 5’ splice-site strength was calculated via Berkeley Drosophila Genome Project’s splice-site predictor for human, with a minimum score threshold of 0.4 (Reese et al. 1997). Branchpoint (BP) prediction and probability scoring were determined using the R package, *branchpointer* (Signal et al. 2017). Capital “A” s designates potential branchpoint adenines, with font size corresponding to BP probability score magnitude (>25% probability was reported), and underlined adenines (<u>A</u>) represent the highest scored BP. Predicted BP with strong relative U2 binding energy scores (>4.0) were marked with an asterisk (*). PPT sequence prediction and scoring were performed as previously described (Corvelo et al. 2010). Sequence prediction (in red) was determined heuristically 5’ to the highest BP score (in red), as well as for maximum length (continued in green). PPT scores were calculated using a custom R script.

### Supplementary Table 2

Endogenous and minigene splice-site strengths were obtained from Berkeley Drosophila Genome Project (BDGP). The acceptor-site of each middle exon corresponds to the 3’ splice-site of the previous adjacent intron and the donor-site of each middle exon corresponds to the 5’ splice-site of the next adjacent intron. *SS: Splice-site strength.

### Supplementary Table 3

Conserved SRSF2 binding-site scores within human *MDM2* exons 3 through 12 were identified using ESEfinder (v2.0) with default threshold of 2.383. Conserved sequences between mouse and human were selected. “Score Average” refers to the arithmetic mean of all conserved bind-site scores; “Score Sum” is the total of all conserved scores; “Score Sum per Exon Length” is the sum of all conserved scores divided by the total number of coding base pairs in the exon.

### Supplementary Table 4

Table of predicted SRSF2 binding-site motifs conserved between human and mouse *MDM2* exons 3 through 12 were identified using ESE-finder (v2.0) with default threshold of 2.383. Motifs with 87.5-100% sequence similarity between human and mouse are in bold, and the scores of motifs with 100% identical motifs are presented as consensus scores. Single nucleotide variants (SNVs) between human and mouse motifs are shown in red.

### Statistical Analyses

Percentages of full-length and exon-excluded products were quantified using ImageQuant TL (Version 8.1). Statistical significance of all results were assessed using the two-tailed Student t-test using GraphPad Prism (Version 8.0) unless otherwise noted.

## Data Availability Statement

All data presented in this manuscript are included in the main and the Supplemental figures. Additional data, materials, or reagents can be requested from the authors via email at.

## Acknowledgments

We would like to thank Juliann Rectenwald and the Ohio State University Comparative Pathology & Mouse Phenotyping Shared Resource for helping with animal necropsies and pathological evaluations. We thank the Chandler lab members for their input on experimental techniques and critical review of this manuscript. This work was funded in part by Pelotonia fellowships to SK and MM, and NIH grant RO1CA262873 to DSC. DSC is also supported by the Cancer Center Core Grant is P30 CA016058 for shared resources.

## Author Contributions

These studies were conceptualized by D.S.C., S.K., M.M. Methodology was designed by D.S.C., S.K., M.M., R.R, Z.L.S, W.D.J, A.G, H.A, C.K. Experiments were performed by S.K., M.M., R.R, Z.L.S, W.D.J, A.G, H.A and C.K. Mutant mice were generated at OSU animal core led by V. C. The data presented here was curated by S.K., M.M., R.R, Z.L.S, W.D.J, A.G, H.A, C.K. and visualized by S.K., M.M., R.R, Z.L.S, W.D.J, A.G, H.A, C.K. The original manuscript was written by S.K., R.R, and Z.L.S, and additional versions were reviewed and edited by D.S.C., S.K. M.M., R.R W.D.J, and Z.L.S. Project supervision and administration were overseen by D.S.C. and funding was acquired by D.S.C.

---

**Supplementary Figure 1:**
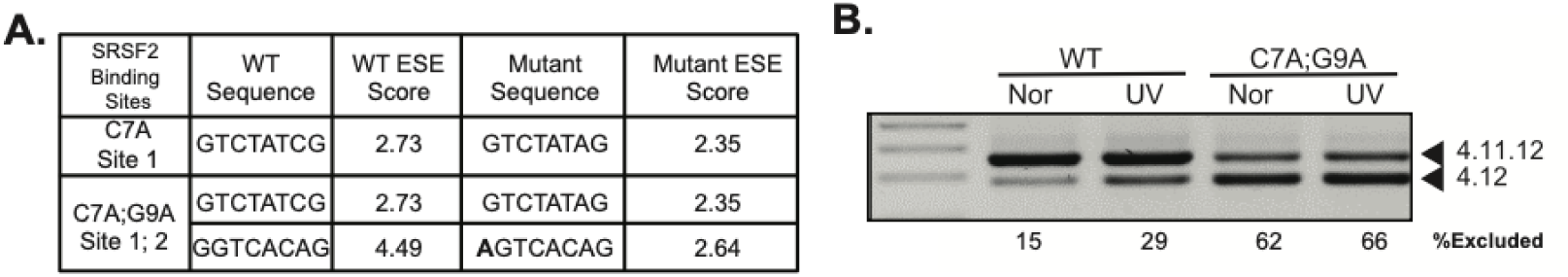
**A.** ESE finder depictions of binding matrix scores of C7A mutant vs C7A;G9A mutant. **B**. RT-PCR of C7A;G9A mutation in mouse *Mdm2* 4-11-12 minigene showed increased exon exclusion compared to wild type minigene even in the absence of genotoxic stress.

**Supplementary Figure 2:**
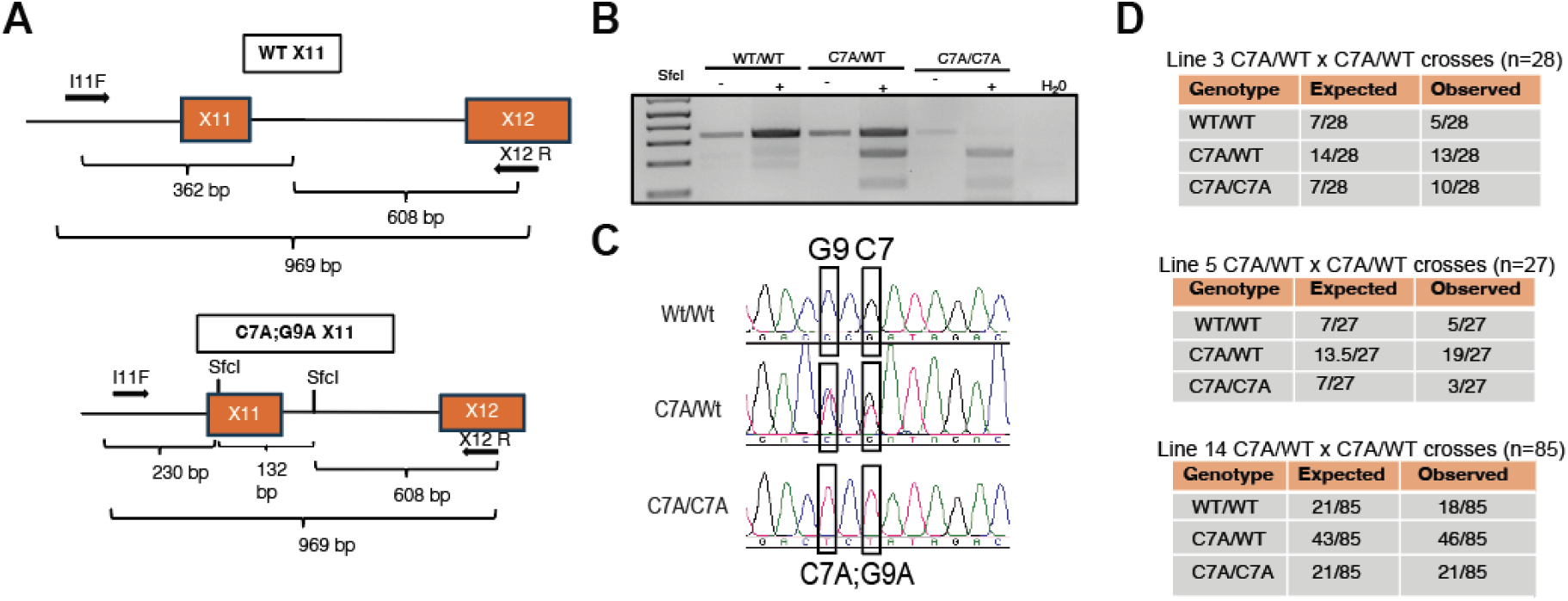
**A.** Schematic depicting the genotyping strategy and represent the primer sequences in *Mdm2* exon 11. The C7A mutation created a restriction digestion-site for the enzyme SfcI. Only mutant cells produce three fragments after digestion with SfcI restriction enzyme. **B**. Genotyping RT-PCR for the CRISPR mice representing the wild type, heterozygous and homozygous mutant mice. **C**. Sanger sequencing to validate the G9A mutation since only C7A mutation can be detected with the restriction digestion. **D.** Table showing number of viable mice from each line demonstrating that the mice were born within mendelian ratio.

**Supplementary Figure 3:**
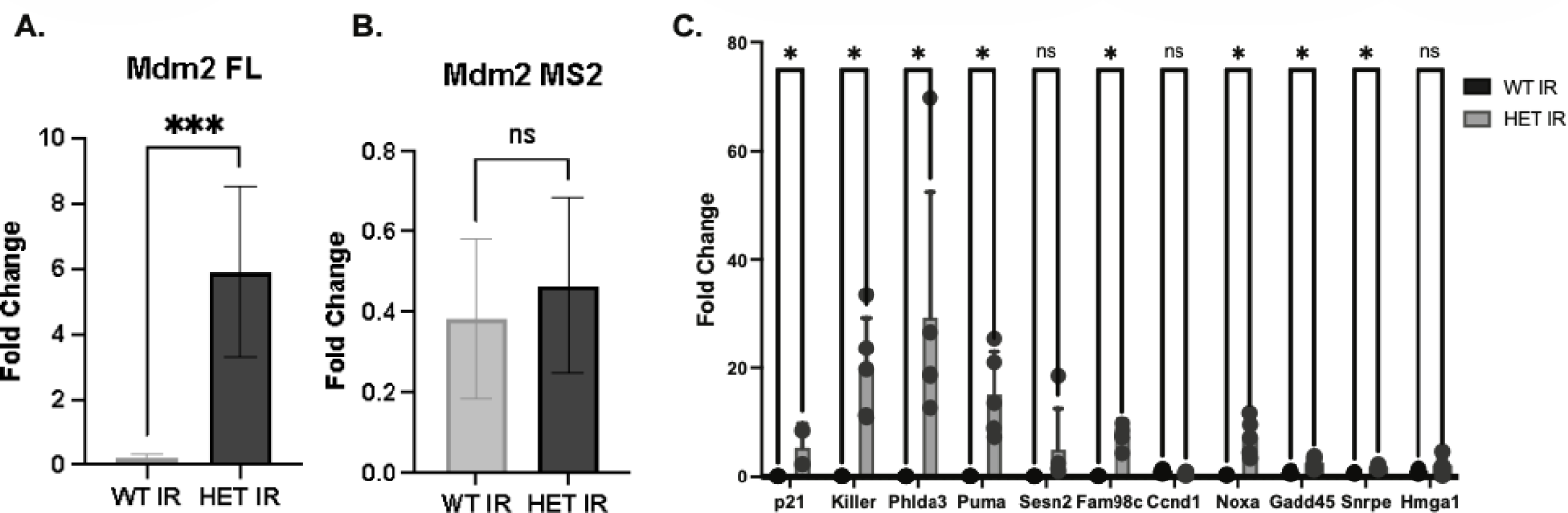
**A, B.** RT-PCR for mouse *Mdm2-FL* and *MS2* in line 3 mice treated with gamma irradiation and harvested after 3 hours post gamma irradiation. **C**. qPCR of p53 target genes for RNA harvested from spleen of WT and C7A;G9A gamma irradiated mice.

**Supplementary Table 1:** Table of predicted SRSF2 binding-site motifs conserved between human and mouse *MDM2* in exons 3 through 12. SRSF2 sequence motifs were identified by ESEfinder (v2.0) with default threshold of 2.383. Motifs with either one single nucleotide variant (shown in red) or 100% sequence similarity between human and mouse are in bold, and motifs that only share exon position between mouse and human are un-bolded in the consensus score column.

| Exon | Length (nt) |  | Percent Sequence Identity | Human |  |  | Mouse |  |  |
| --- | --- | --- | --- | --- | --- | --- | --- | --- | --- |
|  | Human | Mouse |  | Position | Motif | Score | Position | Motif | Score |
| **4** | 75 | 75 | 90.67 | **5** | **GACCAAA**G**** | **3.318484** | **5** | **GACCAAAA** | **2.868588** |
|  |  |  |  | 11 | AGCCATTG | 2.498156 |  |  |  |
|  |  |  |  | **36** | **GTCT**G**TTG** | **2.946445** | **36** | **GTCC**G**TTG** | **3.177286** |
| **5** | 134 | 134 | 85.82 | **16** | **GGCCAGTA** | **4.614038** | **16** | **GGCCAGTA** | **4.614038** |
|  |  |  |  | **83** | **ATCT**T**CTA** | **2.61193** | **83** | **ATCT**C**CTA** | **3.371171** |
|  |  |  |  | 99 | GTTTGGCG | 2.546779 |  |  |  |
|  |  |  |  |  |  |  | 104 | GAGTCCCG | 2.907951 |
|  |  |  |  |  |  |  | 106 | GTCCCGAG | 3.86556 |
|  |  |  |  | **114** | **CTTCTCTG** | **2.447293** | **114** | **TTTCTCTG** | **2.426165** |
| **6** | 50 | 50 | 78.00 | **16** | **GATCTACA** | **2.391501** | **16** | **GATCTACA** | **2.391501** |
| **7** | 68 | 59 | 81.36 | 8 | GGACTCAG | 3.658624 |  |  |  |
|  |  |  |  |  |  |  | 10 | CATCGCTG | 2.595681 |
| **8** | 97 | 97 | 82.47 | 14 | AGCTTCAG | 2.686393 | 14 | CGCCACCA | 2.856393 |
|  |  |  |  |  |  |  | 36 | ATCTTCTG | 3.061826 |
|  |  |  |  | 48 | GGTTTCTA | 4.442748 |  |  |  |
| **9** | 161 | 164 | 67.11 |  |  |  | 5 | GAACACAG | 3.584598 |
|  |  |  |  | 42 | CGCCACAA | 2.477722 | 42 | CGCCGCAG | 2.502354 |
|  |  |  |  |  |  |  | 49 | GGTCCCTG | 5.882726 |
|  |  |  |  |  |  |  | 56 | GTCCTTTG | 3.315724 |
|  |  |  |  |  |  |  | 63 | GATCCGAG | 3.583454 |
|  |  |  |  |  |  |  | 75 | GGTCTGTG | 4.208176 |
|  |  |  |  |  |  |  | 140 | GTCCACAG | 4.308454 |
|  |  |  |  | 145 | GGACGCCA | 3.448961 |  |  |  |
| **10** | 156 | 156 | 87.18 |  |  |  | 4 | GCATTCTG | 2.414814 |
|  |  |  |  | **48** | **GGATT**C**AG** | **3.427783** | **48** | **GGATT**C**AG** | **3.427783** |
|  |  |  |  |  |  |  | 54 | AGTTTCTG | 3.140572 |
|  |  |  |  | **66** | **GTTT**A****G**TG** | **3.616483** | **66** | **GTTT**A****G****C**G** | **2.972043** |
|  |  |  |  |  |  |  | 89 | AGTCTCTG | 3.371413 |
|  |  |  |  |  |  |  | 106 | GATTACAG | 3.795507 |
|  |  |  |  |  |  |  | 147 | GGATGATG | 3.03566 |
| **11** | 78 | 78 | 82.05 |  |  |  | 1 | GTCTATCG | 2.727269 |
|  |  |  |  | 9 | AGTTACTG | 3.427398 | 9 | GGTCACAG | 4.3872 |
|  |  |  |  | **57** | **AGAT**C**CTG** | **3.458063** | **57** | **AGAT**C**CTG** | **3.458063** |

**Supplementary Table 2:**
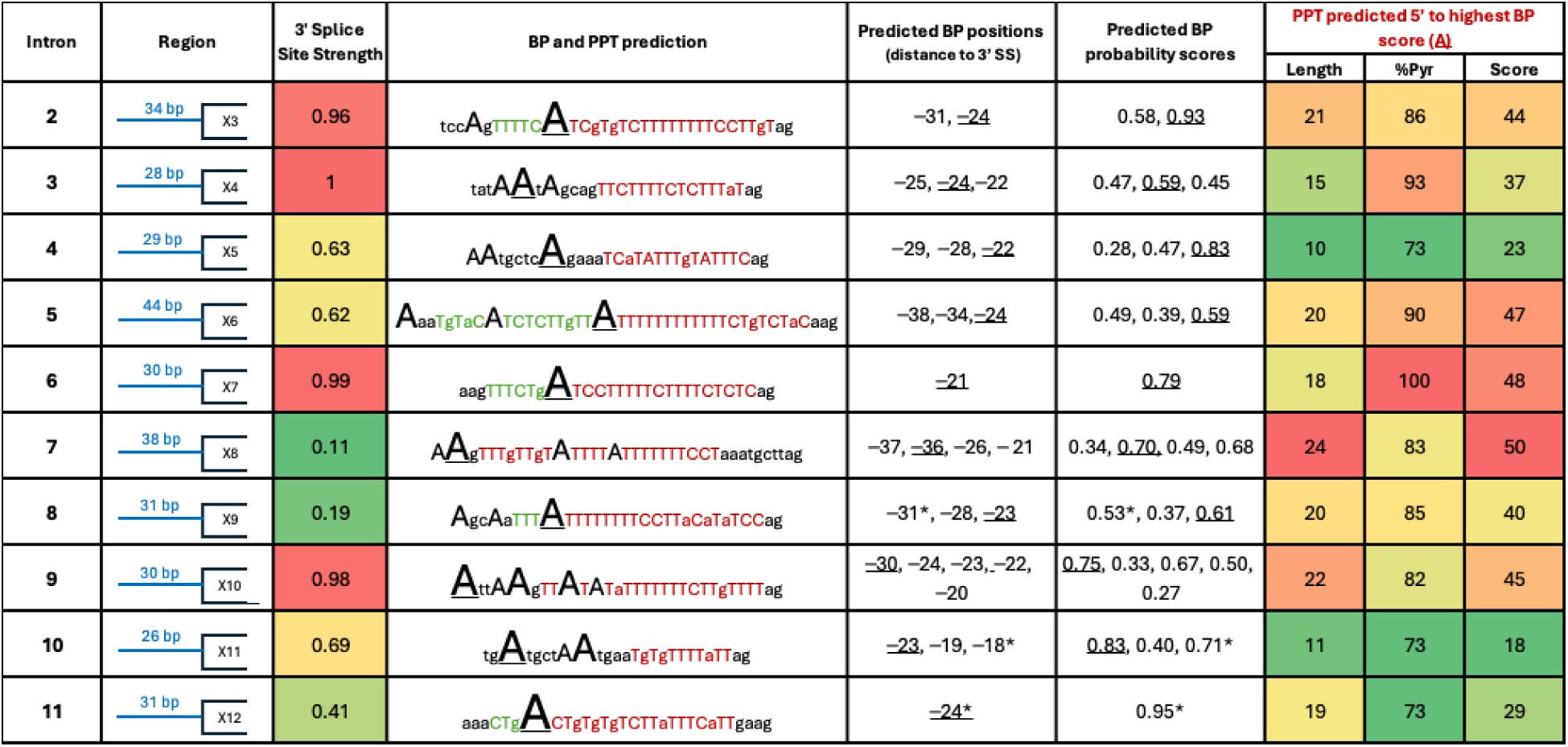
Table of bioinformatically predicted 3’ splice-site (SS) strength, branch point (BP) position and probability score and polypyridine tract (PPT) length, percent, and score for *MDM2* introns 3 through 11. 5’ splice-site strength was calculated via Berkeley Drosophila Genome Project’s splice-site predictor for human; minimum score threshold was 0.4 (Reese MG, et al., 1997). BP prediction and probability scoring was determined using the R package, *branchpointer* (Signal, et al., 2018). Capital “A”s designate potential branchpoint adenines with font size corresponding to BP probability score magnitude (>25% probability were reported), and underlined adenines (<u>A</u>) represent the highest scored BP. Strong relative U2 binding energy scores (>4.0) were noted for the predicted BPs (*). PPT sequence prediction and scoring were described previously in Corvelo, et al., 2010. Sequence prediction (in red) was determined heuristically 5’ to the highest BP score (in red), as well as for maximum length (continued in green). PPT scores were calculated using a custom R script.

**Supplementary Table 3:** Table showing the difference in splice-site strength among the minigene constructs designed for figure 1 (Minigene Panel-I) and 2 (Minigene Panel-II). In this table, the acceptor-site of each intervening exon is the 3’ splice-site of the previous adjacent intron and the donor-site of each intervening exon is the 5’ splice-site of the next adjacent intron. *SS: Splice-sitestrength.

| Exon | Minigene Fig.1 SS Strength |  | Endogenous + Minigene Fig. 2 SS Strength |  |
| --- | --- | --- | --- | --- |
|  | Acceptor Site | Donor Site | Acceptor Site | Donor Site |
| 4 | 0.96 | 0.96 | 1 | 1 |
| 5 | 0.66 | 0.98 | 0.63 | 0.98 |
| 6 | - | 1 | 0.62 | 1 |
| 7 | 0.78 | 0.89 | 0.99 | 0.89 |
| 8 | - | 0.92 | 0.11 | 0.92 |
| 9 | - | 0.60 | 0.19 | 0.60 |
| 10 | 0.77 | 0.34 | 0.98 | 0.34 |
| 11 | 0.69 | 0.98 | 0.69 | 0.96 |

**Supplementary Table 4:** Conserved SRSF2 binding-site score within human *MDM2* exons 4 through 11. SRSF2 sequence motifs were identified by ESEfinder (v2.0) with default threshold of 2.383; conserved sequences between mouse and human were selected. “Score Average” = arithmetic mean of all conserved bind-site scores; “Score Sum” = sum of all conserved scores; “Score Sum per Exon Length” = the sum of all conserved scores divided by the total number of nucleotides in each exon.

| Exon | Score Average | Score Sum | Score Sum per Exon Length |
| --- | --- | --- | --- |
| 4 | 3.1324645 | 6.264929 | 0.083532387 |
| 5 | 3.217377667 | 9.652133 | 0.072030843 |
| 6 | 2.391501 | 2.391501 | 0.04783002 |
| 7 | 0 | 0 | 0 |
| 8 | 2.686393 | 2.686393 | 0.027694773 |
| 9 | 2.477722 | 2.477722 | 0.015389578 |
| 10 | 3.522133 | 7.044266 | 0.045155551 |
| 11 | 3.4427305 | 6.885461 | 0.088275141 |

**Supplementary Table 5:**
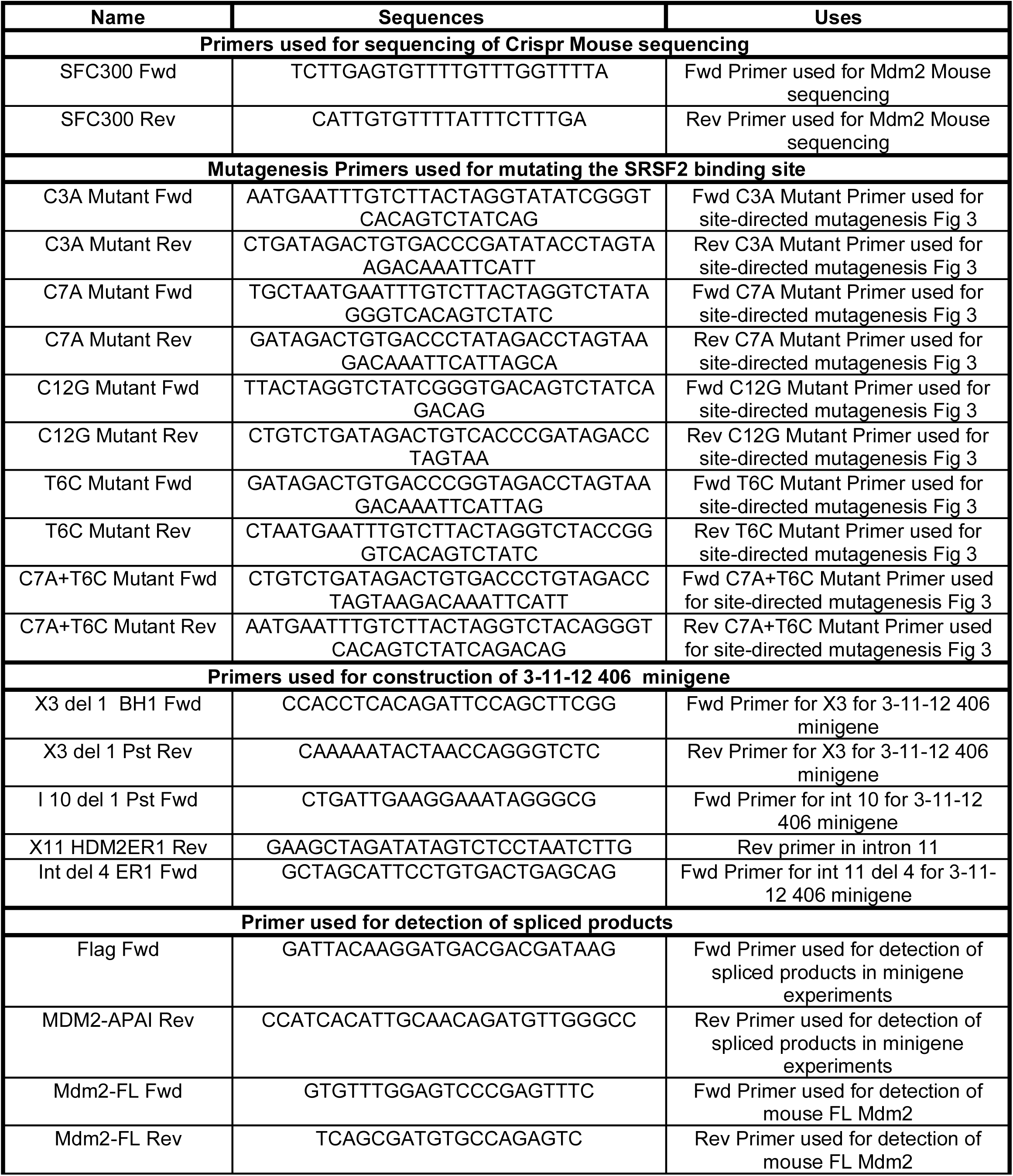
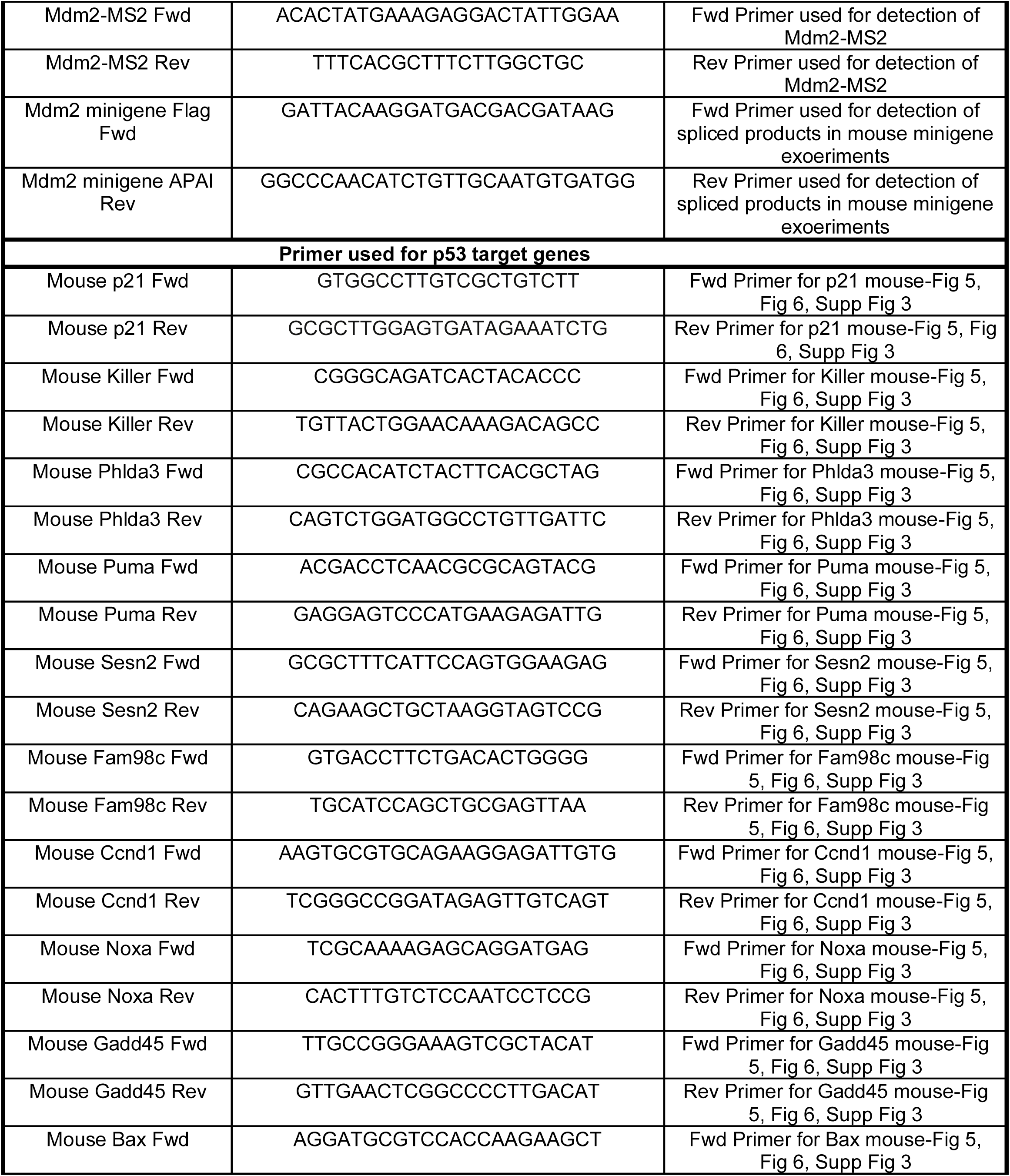
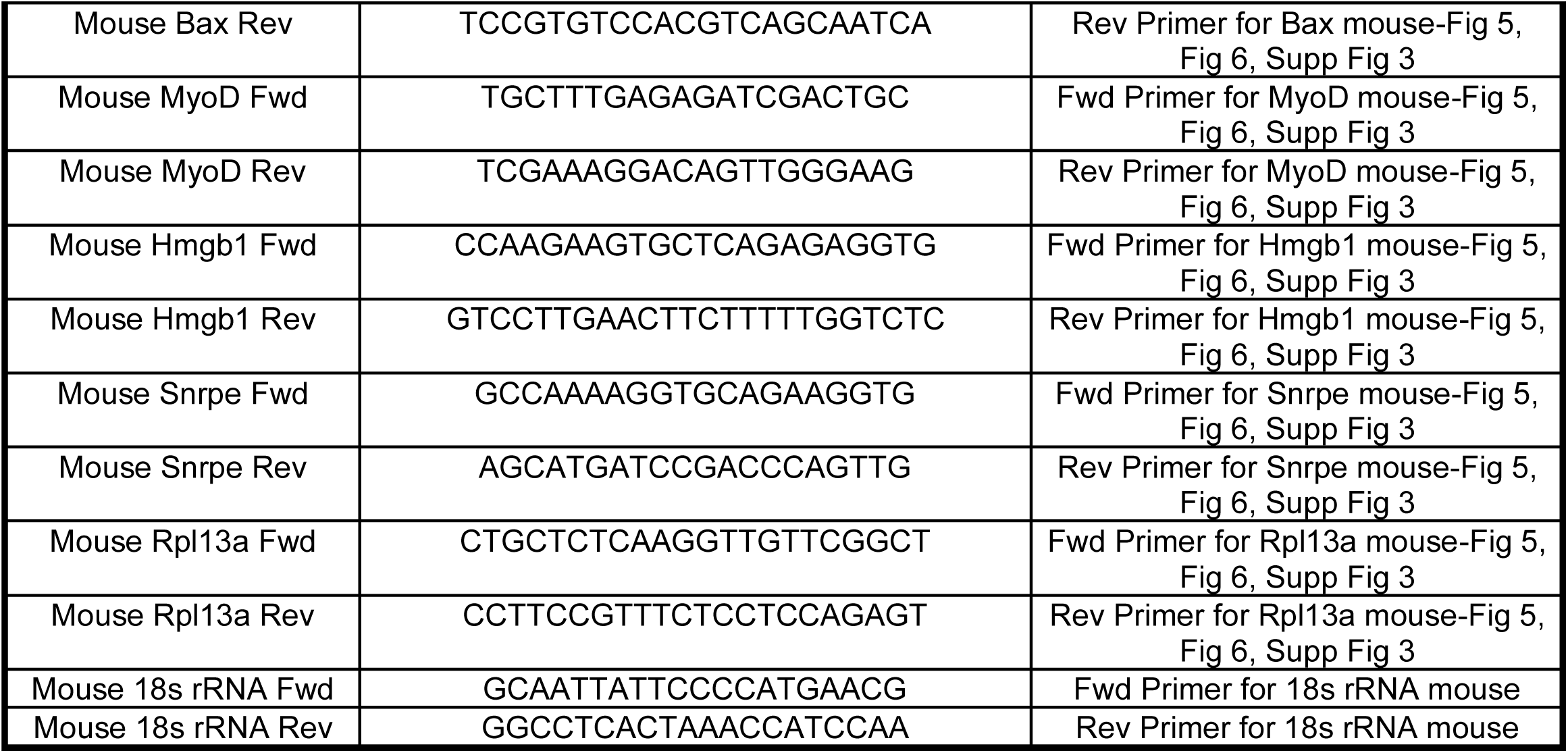
Primers used throughout the paper.

## References

1. Baralle FE, Giudice J. Alternative splicing as a regulator of development and tissue identity. Nat Rev Mol Cell Biol. 2017 Jul;18(7):437–51.

2. Bradley RK, Anczuków O. RNA splicing dysregulation and the hallmarks of cancer. Nat Rev Cancer. 2023 Mar;23(3):135–55.

3. Dvinge H, Kim E, Abdel-Wahab O, Bradley RK. RNA splicing factors as oncoproteins and tumour suppressors. Nat Rev Cancer. 2016 Jul;16(7):413–30.

4. Oltean S, Bates DO. Hallmarks of alternative splicing in cancer. Oncogene. 2014 Nov 13;33(46):5311–8.

5. Wang Y, Liu J, Huang BO, Xu YM, Li J, Huang LF, et al. Mechanism of alternative splicing and its regulation. Biomed Rep. 2015 Mar;3(2):152–8.

6. Cartegni L, Wang J, Zhu Z, Zhang MQ, Krainer AR. ESEfinder: A web resource to identify exonic splicing enhancers. Nucleic Acids Res. 2003 Jul 1;31(13):3568–71.

7. Chandler DS, Singh RK, Caldwell LC, Bitler JL, Lozano G. Genotoxic stress induces coordinately regulated alternative splicing of the p53 modulators MDM2 and MDM4. Cancer Res. 2006 Oct 1;66(19):9502–8.

8. Biamonti G, Caceres JF. Cellular stress and RNA splicing. Trends Biochem Sci. 2009 Mar;34(3):146–53.

9. Bonnal SC, López-Oreja I, Valcárcel J. Roles and mechanisms of alternative splicing in cancer - implications for care. Nat Rev Clin Oncol. 2020 Aug;17(8):457–74.

10. Dreyfuss G, Matunis MJ, Piñol-Roma S, Burd CG. hnRNP proteins and the biogenesis of mRNA. Annu Rev Biochem. 1993;62:289–321.

11. Bradley T, Cook ME, Blanchette M. SR proteins control a complex network of RNA-processing events. RNA N Y N. 2015 Jan;21(1):75–92.

12. Venables JP, Koh CS, Froehlich U, Lapointe E, Couture S, Inkel L, et al. Multiple and specific mRNA processing targets for the major human hnRNP proteins. Mol Cell Biol. 2008 Oct;28(19):6033–43.

13. Comiskey DF, Jacob AG, Singh RK, Tapia-Santos AS, Chandler DS. Splicing factor SRSF1 negatively regulates alternative splicing of MDM2 under damage. Nucleic Acids Res. 2015 Apr 30;43(8):4202–18.

14. Fu XD, Maniatis T. The 35-kDa mammalian splicing factor SC35 mediates specific interactions between U1 and U2 small nuclear ribonucleoprotein particles at the 3’ splice site. Proc Natl Acad Sci U S A. 1992 Mar 1;89(5):1725–9.

15. Fu XD, Mayeda A, Maniatis T, Krainer AR. General splicing factors SF2 and SC35 have equivalent activities in vitro, and both affect alternative 5’ and 3’ splice site selection. Proc Natl Acad Sci U S A. 1992 Dec 1;89(23):11224–8.

16. Liu S, Zhou B, Wu L, Sun Y, Chen J, Liu S. Single-cell differential splicing analysis reveals high heterogeneity of liver tumor-infiltrating T cells. Sci Rep. 2021 Mar 5;11:5325.

17. Qian W, Liang H, Shi J, Jin N, Grundke-Iqbal I, Iqbal K, et al. Regulation of the alternative splicing of tau exon 10 by SC35 and Dyrk1A. Nucleic Acids Res. 2011 Aug;39(14):6161–71.

18. Loh TJ, Moon H, Cho S, Jung DW, Hong SE, Kim DH, et al. SC35 promotes splicing of the C5-V6-C6 isoform of CD44 pre-mRNA. Oncol Rep. 2014 Jan;31(1):273–9.

19. Marine JC, Lozano G. Mdm2-mediated ubiquitylation: p53 and beyond. Cell Death Differ. 2010 Jan;17(1):93–102.

20. Shi D, Gu W. Dual Roles of MDM2 in the Regulation of p53: Ubiquitination Dependent and Ubiquitination Independent Mechanisms of MDM2 Repression of p53 Activity. Genes Cancer. 2012 Mar;3(3–4):240–8.

21. Nag S, Qin J, Srivenugopal KS, Wang M, Zhang R. The MDM2-p53 pathway revisited. J Biomed Res. 2013 Jul;27(4):254–71.

22. Sigalas I, Calvert AH, Anderson JJ, Neal DE, Lunec J. Alternatively spliced mdm2 transcripts with loss of p53 binding domain sequences: transforming ability and frequent detection in human cancer. Nat Med. 1996 Aug;2(8):912–7.

23. Singh RK, Tapia-Santos A, Bebee TW, Chandler DS. Conserved sequences in the final intron of MDM2 are essential for the regulation of alternative splicing of MDM2 in response to stress. Exp Cell Res. 2009 Nov 15;315(19):3419–32.

24. Comiskey DF, Montes M, Khurshid S, Singh RK, Chandler DS. SRSF2 Regulation of MDM2 Reveals Splicing as a Therapeutic Vulnerability of the p53 Pathway. Mol Cancer Res MCR. 2020 Feb;18(2):194–203.

25. Jacob AG, O’Brien D, Singh RK, Comiskey DF, Littleton RM, Mohammad F, et al. Stress-induced isoforms of MDM2 and MDM4 correlate with high-grade disease and an altered splicing network in pediatric rhabdomyosarcoma. Neoplasia N Y N. 2013 Sep;15(9):1049–63.

26. Huun J, Gansmo LB, Mannsåker B, Iversen GT, Øvrebø JI, Lønning PE, et al. Impact of the MDM2 splice-variants MDM2-A, MDM2-B and MDM2-C on cytotoxic stress response in breast cancer cells. BMC Cell Biol. 2017 Apr 17;18(1):17.

27. de Faria FCC, Khurshid S, Sarchet P, Tahara S, Casadei L, Grignol V, et al. Oncogenic Functions of Alternatively Spliced MDM2-ALT2 Isoform in Retroperitoneal Liposarcoma. Int J Mol Sci. 2024 Dec 17;25(24):13516.

28. Corvelo A, Hallegger M, Smith CWJ, Eyras E. Genome-wide association between branch point properties and alternative splicing. PLoS Comput Biol. 2010 Nov 24;6(11):e1001016.

29. Jacob AG, Singh RK, Mohammad F, Bebee TW, Chandler DS. The splicing factor FUBP1 is required for the efficient splicing of oncogene MDM2 pre-mRNA. J Biol Chem. 2014 Jun 20;289(25):17350–64.

30. Reese MG, Eeckman FH, Kulp D, Haussler D. Improved splice site detection in Genie. J Comput Biol J Comput Mol Cell Biol. 1997;4(3):311–23.

31. Taheri M, Abdullah SR, Saber AF, Samsami M, Hussen BM. The dual role of cyclin D1: Unraveling its tumor-promoting mechanisms and opportunities for therapeutics. Int J Biol Macromol. 2025 Dec 1;333:148957.

32. Wang J, Su W, Zhang T, Zhang S, Lei H, Ma F, et al. Aberrant Cyclin D1 splicing in cancer: from molecular mechanism to therapeutic modulation. Cell Death Dis. 2023 Apr 6;14(4):244.

33. Comiskey DF, Jacob AG, Sanford BL, Montes M, Goodwin AK, Steiner H, et al. A novel mouse model of rhabdomyosarcoma underscores the dichotomy of MDM2-ALT1 function in vivo. Oncogene. 2018 Jan 4;37(1):95–106.

34. Pant V, Xiong S, Jackson JG, Post SM, Abbas HA, Quintás-Cardama A, et al. The p53-Mdm2 feedback loop protects against DNA damage by inhibiting p53 activity but is dispensable for p53 stability, development, and longevity. Genes Dev. 2013 Sep 1;27(17):1857–67.

35. Bergfort A, Gordon JM, Gazzara MR, Hung CT, Lee B, Barash Y, et al. The exon junction complex coordinates the cotranscriptional inclusion of blocks of neighboring exons. Genes Dev. 2026 Jan 5;40(1–2):94–109.

36. Shine M, Gordon J, Schärfen L, Zigackova D, Herzel L, Neugebauer KM. Co-transcriptional gene regulation in eukaryotes and prokaryotes. Nat Rev Mol Cell Biol. 2024 Jul;25(7):534–54.

37. Choquet K, Baxter-Koenigs AR, Dülk SL, Smalec BM, Rouskin S, Churchman LS. Pre-mRNA splicing order is predetermined and maintains splicing fidelity across multi-intronic transcripts. Nat Struct Mol Biol. 2023 Aug;30(8):1064–76.

38. Wan Y, Anastasakis DG, Rodriguez J, Palangat M, Gudla P, Zaki G, et al. Dynamic imaging of nascent RNA reveals general principles of transcription dynamics and stochastic splice site selection. Cell. 2021 May 27;184(11):2878–2895.e20.

39. Donovan BT, Wang B, Garcia GR, Mount SM, Larson DR. REAL-TIME VISUALIZATION OF SPLICEOSOME ASSEMBLY REVEALS BASIC PRINCIPLES OF SPLICE SITE SELECTION. bioRxiv. 2024 Jul 13;2024.07.12.603320.

40. Glidden DT, Buerer JL, Saueressig CF, Fairbrother WG. Hotspot exons are common targets of splicing perturbations. Nat Commun. 2021 May 12;12(1):2756.

41. Kim SW, Taggart AJ, Heintzelman C, Cygan KJ, Hull CG, Wang J, et al. Widespread intra-dependencies in the removal of introns from human transcripts. Nucleic Acids Res. 2017 Sep 19;45(16):9503–13.

42. García-Ruiz S, Zhang D, Gustavsson EK, Rocamora-Perez G, Grant-Peters M, Fairbrother-Browne A, et al. Splicing accuracy varies across human introns, tissues, age and disease. Nat Commun. 2025 Jan 27;16(1):1068.

43. Gohr A, Torres-Méndez A, Bonnal S, Irimia M. Insplico: Effective computational tool for studying intron splicing order genome-wide with short and long RNA-seq reads [Internet]. bioRxiv; 2022 [cited 2026 Feb 10]. p. 2022.08.15.503947. Available from: https://www.biorxiv.org/content/10.1101/2022.08.15.503947v1

44. De Conti L, Baralle M, Buratti E. Exon and intron definition in pre-mRNA splicing. Wiley Interdiscip Rev RNA. 2013;4(1):49–60.

45. Rogalska ME, Vivori C, Valcárcel J. Regulation of pre-mRNA splicing: roles in physiology and disease, and therapeutic prospects. Nat Rev Genet. 2023 Apr;24(4):251–69.

46. Zhang Y, Qian J, Gu C, Yang Y. Alternative splicing and cancer: a systematic review. Signal Transduct Target Ther. 2021 Feb 24;6(1):78.

47. El Marabti E, Younis I. The Cancer Spliceome: Reprograming of Alternative Splicing in Cancer. Front Mol Biosci. 2018;5:80.

48. Stanley RF, Abdel-Wahab O. Dysregulation and therapeutic targeting of RNA splicing in cancer. Nat Cancer. 2022 May;3(5):536–46.

49. Zeeshan S, Dalal B, Arauz RF, Zingone A, Harris CC, Khiabanian H, et al. Global profiling of alternative splicing in non-small cell lung cancer reveals novel histological and population differences. Oncogene. 2025 Apr;44(14):958–67.

50. Urbanski LM, Leclair N, Anczuków O. Alternative-splicing defects in cancer: Splicing regulators and their downstream targets, guiding the way to novel cancer therapeutics. Wiley Interdiscip Rev RNA. 2018 Jul;9(4):e1476.

51. Khurshid S, Montes M, Comiskey DF, Shane B, Matsa E, Jung F, et al. Splice-switching of the insulin receptor pre-mRNA alleviates tumorigenic hallmarks in rhabdomyosarcoma. NPJ Precis Oncol. 2022 Jan 11;6(1):1.

52. Chen H, Tang J, Xiang J. Alternative Splicing in Tumorigenesis and Cancer Therapy. Biomolecules. 2025 May 29;15(6):789.

